# Connectomic reconstruction from hippocampal CA3 reveals spatially graded mossy fiber inputs and selective feedforward inhibition to pyramidal cells

**DOI:** 10.1101/2025.07.09.663979

**Authors:** Zhihao Zheng, Changjoo Park, Eric W. Hammerschmith, Ran Lu, Szi-Chieh Yu, Marissa Sorek, Ben Silverman, Chris S. Jordan, Amy R. Sterling, William M. Silversmith, Forrest Collman, H. Sebastian Seung, David W. Tank

**Affiliations:** Princeton Neuroscience Institute, Princeton University, Princeton, NJ, USA; Allen Institute for Brain Science, Seattle, WA, USA; Computer Science Department, Princeton University, Princeton, NJ, USA; Bezos Center for Neural Circuit Dynamics, Princeton University, Princeton, NJ, USA; Department of Molecular Biology, Princeton University, Princeton, NJ, USA

## Abstract

The mossy fiber (MF) connections to pyramidal cells in hippocampal CA3 are hypothesized to participate in pattern separation and memory encoding, yet no large-scale neuronal wiring diagram exists for these connections. We assembled a 3D electron microscopy volume (∼1×1×0.1mm^3^) from mouse hippocampal CA3. By proofreading and automated segmentation, we reconstructed and classified all soma-containing neurons—including 1,815 pyramidal cells and 229 inhibitory cells—and over 55,000 MFs. Pyramidal cells receive more numerous MF inputs along a proximodistal gradient. Some distal cells show surprisingly high convergence via relatively small terminals with fewer vesicles. Pyramidal cells share significantly more MF inputs than networks randomized by degree-preserving swap, and are better approximated by networks randomized by proximity-preserving swap. We identify a feedforward inhibitory circuit from MFs via perisomatic interneurons that selectively target a pyramidal subtype. We demonstrated large-scale mapping across levels in the hippocampus—from circuits to cell types to vesicles. The dataset is shared through Pyr, an online platform for hippocampal connectomics.

## Introduction

The hippocampus has been implicated in episodic and spatial memory. Synaptic plasticity in the hippocampus is widely considered to be a cellular mechanism of learning and memory. To study the structural correlates of this plasticity, serial electron microscopy (EM) has been used to characterize hippocampal synapses in 3D^1–7^. These studies could be performed with relatively small EM volumes. Larger volumes are required to study the connectivity of circuits in the context of the morphology and identity of the constituent neurons because synapses are distributed along axons and dendrites that span millimeters in scale. While the sizes of 3D EM volumes for smaller model organisms and cortex have increased significantly^8–12^, the scale-up of synapse-resolution volumetric imaging and reconstruction for the hippocampus is still in its infancy.

Recent studies have attempted to increase the size of EM volumes in the rodent hippocampus. To study the distribution of putative entorhinal inputs, conventional TEM imaging of serial ultrathin sections was used to acquire a volume that spans two sublayers of CA1^13^ (350 x 200 × 17 μm^3^). Sammons et al.^14^ acquired a volume that encompasses all layers of CA3 with 1 mm^2^ sections (965 × 808 × 62 μm^3^). Analysis of pyramidal cell connections in the dataset has revealed a high recurrent connectivity rate. In parallel, a larger EM volume of CA1 (1.1 x 0.92 x 0.143 mm³) was recently reported, along with analysis of synaptic weights across a large number of synapses^15,16^. These volumes are sparsely reconstructed and the analysis is primarily focused on recurrent connectivity. Here, we focus on a more complete reconstruction in the CA3 region of the mouse hippocampus, with an emphasis on mossy fiber (MF) and inhibitory inputs to pyramidal cells.

The hippocampus, classically known as the archicortex, is organized as a single layer of principal cells among a diverse array of interneurons, across its three major regions: dentate gyrus (DG), CA3, and CA1. Granule cells in the DG give rise to MFs, which provide a major excitatory input pathway to pyramidal cells of the CA3 region. The DG is often modeled as a circuit with the computational role of pattern separation^17–19^, while CA3, due to its extensive recurrent collaterals, is conjectured to support pattern completion^20–22^. However, morphological heterogeneity of the single-layer CA3 pyramidal cells has been documented early on and is increasingly associated with physiological and functional heterogeneity. Along the proximodistal axis, a gradient in the properties of the CA3 pyramidal cells have been characterized in a number of anatomical, physiological, functional, and behavioral studies^23–27^. Along the superficial-deep (radial) axis, pyramidal cells are organized into two sublayers, distinguished by the abundance of MF inputs, with distinct physiological properties and proposed functions^28–31^. In this study, we aim to characterize synaptic ultrastructural correlates of these systematic variations of CA3 pyramidal cells.

Mapping the MF to CA3 pyramidal connectivity will inform models of how a pattern separation circuit is transformed into a pattern completion circuit in the brain^32,33^. The MF boutons are typically large enough to be seen with light microscopy, and prior studies have quantified the number and spatial distribution of boutons on MFs^34–36^. However, current understanding of MF–CA3 connectivity relies primarily on these bouton counts along with estimates of the number of CA3 pyramidal neurons, under the assumption that each presynaptic bouton contacts only a single pyramidal cell^1^. While MF terminals are typically large, they exhibit substantial morphological variability, with some considerably smaller^37,38^. The one-bouton-one-CA3-cell assumption remains untested at scale. Furthermore, given the heterogeneity of CA3 pyramidal cells, it is important to study spatial bias of their MF inputs, particularly along the proximodistal axis. While MF-CA3 connectivity has typically been modeled as uniformly random, detailed mapping that captures the heterogeneity of these connections could provide a starting point to think about how connectivity variations contribute to computational functions such as pattern separation and pattern completion.

We have acquired, assembled, and automatically segmented a 0.1 mm^3^ volume of hippocampal CA3, including neuronal and non-neuronal cells, cell nuclei, and synapses. Dense segmentation is hosted on a Connectome Annotation Versioning Engine (CAVE) architecture for real-time collaborative proofreading, annotation, and analysis^39^. Additional segmentation of subcellular structures such as boutons and vesicles is demonstrated. We proofread and classified all soma-containing neurons and ∼55,000 MFs in the volume. With these reconstructions we have characterized three features of connectivity onto CA3 pyramidal cells: (1) spatial gradients, with proximodistal increases in MF inputs; (2) high convergence, with a small subset of pyramidal cells receiving a surprisingly large number of MF inputs; (3) cell-type specificity, with perisomatic inhibitory axons targeting distinct subtypes of pyramidal cells. These findings constrain computational models of hippocampal circuits, and exemplify the utility of the dataset. We are releasing the dataset as a resource to the community and we expect other researchers will find it useful for addressing their own questions.

## Results

### Dataset overview

A tissue block from the dorsal hippocampus of an adult female mouse (C57BL/6J, 4 months old) was stained using an established protocol^40^ and imaged with X-ray μCT to locate the CA3 region (Fig. 1a-b). The CA3 containing region of the block was sectioned and imaged using our previously reported connectomic imaging pipeline featuring beam deflection transmission electron microscopy (TEM)^41^ and GridTape technology^10^. As the first large-scale acquisition utilizing this pipeline, the CA3 volume spans 1 x 1 x 0.092 mm^3^ at an imaging voxel size of 3 × 3 × 45 nm (Fig. 1c). The volume comprises 2,048 sections cut at 45 nm along the z-axis (z-layer = 96 to 2143), including 39 missing sections (Extended Data Fig. 1a). The largest gap spans five consecutive sections (z-layer = 1262–1266; Extended Data Fig. 1b-c).

**Fig. 1:**
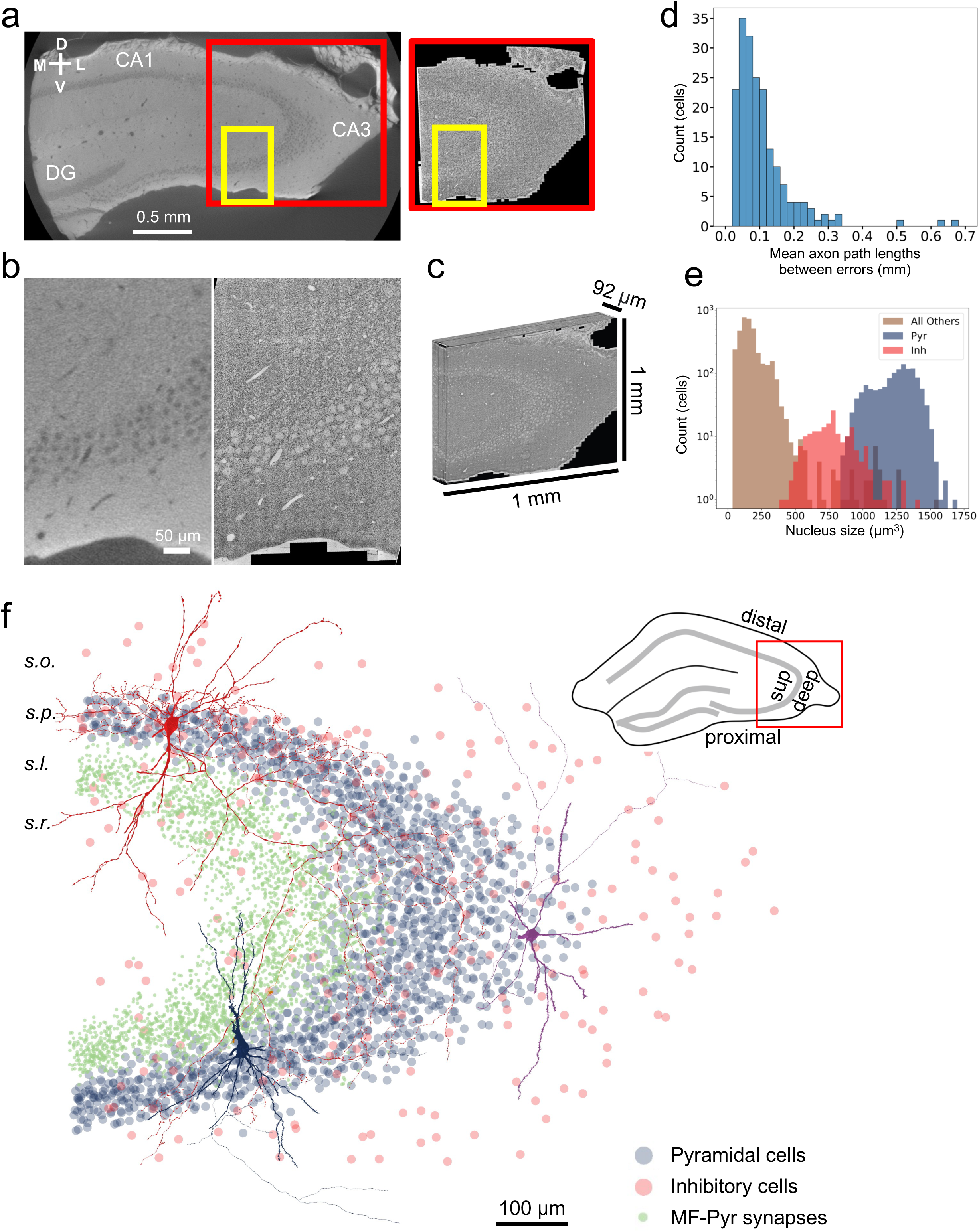
Electron microscopy reconstruction from a hippocampus CA3 volume. **a,** An EM-stained block of hippocampal tissue is first imaged in µCT (left) and sectioned for EM imaging (right). Images of a thin section acquired at 3 nm/pix are stitched together into a montage (right square, ∼1 mm^2^). **b,** Zoom-in (yellow rectangle in **a**) shows the same area where the µCT and EM images were registered. **c,** Dimensions of the 3D EM dataset of CA3 totalling 0.1 mm^3^. **d**, Histogram of mean axon lengths between errors for pyramidal cells with total axon length longer than 0.5 mm (mean 108 µm). **e,** Nucleus size distribution of pyramidal, inhibitory, and other cells from automated nucleus segmentation. **f,** Spatial distribution of pyramidal and inhibitory cells along with mossy fiber–pyramidal (MF-Pyr) synapses. When size-thresholded, the largest synapses—MF-Pyr synapses—occupy a specific layer (s.l.). Three reconstructed cells are shown: a thorny pyramidal cell (dark blue), a sparsely thorny pyramidal cell (purple), and a perisomatic inhibitory cell (red). The inset (upper right) depicts transverse (proximodistal) and radial (superficial-deep) axes of CA3. sup, superficial; s.o., stratum oriens; s.p., stratum pyramidale; s.l., stratum lucidum, s.r., stratum radiatum.

The EM images were aligned with deep learning techniques^42^. Prior algorithms and software for segmentation^43^ were modified to implement a multi-resolution approach^44^, which reduced merge errors between large objects (Methods). The segmented volume contains 4 trillion voxels at a voxel size of 18 × 18 × 45 nm³. We assessed segmentation quality by examining thin axons and narrow spine necks, two challenging structures for automated reconstruction. Analysis of pyramidal axons longer than 0.5 mm shows an average axon length of 108 μm between errors (Fig. 1d; Methods). Assessments of spine attachments show 2 merge errors and 23 false negatives out of 441 ground-truth spines from a pyramidal cell (Extended Data Fig. 2a). Despite the thicker sections (45 nm), our segmentation accuracy is highly competitive with other contemporary mammalian connectomic datasets^8,11,12^. As result, the mean proofreading rate for pyramidal axons is 11.3 mm/hr (Extended Data Fig. 2b-c), several times faster than previous estimates^8,12^. The leading causes of segmentation errors are consecutive missing sections (Methods; Extended Data Fig. 2d-f). We exhaustively proofread two select regions spanning the largest gap, and all segments—including fine processes—could either be traced through the gap (> 95%) or terminated (Extended Data Fig. 1b-c). We concluded that the proofread segmentation is highly accurate in spite of the missing sections.

We have proofread and classified all neurons with somata within the volume—1,815 pyramidal cells and 229 inhibitory neurons(Fig. 1f; Extended Data Fig. 3a-b)—as well as over 55,000 MF axons and 91 inhibitory axons lacking somata but with substantial axonal arbors within the volume. The proofread segments account for 40% of all voxels (Extended Data Fig. 2g). We have developed a website (<u>Pyr.ai</u>, Extended Data Fig. 9) as the main portal for accessing the dataset, providing EM imagery, reconstructions, and cell type classifications for download, programmatic access, and interactive browsing.

The remaining un-proofread segments, while vast in number, are generally smaller in size. Nucleus size distributions show that neurons have larger nuclei than non-neuronal cells, with pyramidal cells larger than inhibitory neurons (Fig. 1e). Inhibitory cells are further classified based on the layer in which the majority of their output synapses are located (Extended Data Fig. 3c). A hallmark of the CA3 region is the large synapses between MF boutons and CA3 pyramidal dendrites (MF-Pyr synapses), located in the soma-free stratum lucidum layer^45^. Indeed, filtering by size reveals that the largest synapses in the dataset are localized to this discrete layer above stratum pyramidale (Fig. 1f).

The volume spans a millimeter in the XY plane along the transverse axis, making it well-suited for study of systematic variations along this dimension. However, its range along the Z axis (92 μm) is limited by number of sections and the sectioning plane was oriented at a slightly acute angle relative to the apical dendrites of pyramidal cells. As a result, neurons at different depths within the volume are truncated at different cellular compartments such as apical dendrites, soma, or basal dendrites (Extended Data Fig. 3d). The volume spans stratum radiatum, stratum lucidum, stratum pyramidale, and stratum oriens, from the superficial to the deep layers (Fig. 1f).

### Mapping MF inputs to CA3 pyramidal cells

The MF-Pyr synapses are physiologically unique^46–49^ and morphologically highly complex: each MF bouton typically envelops a cluster of densely branched dendritic spines—thorny excrescences (TEs)—located on the proximal apical dendrites^1,5–7,50,51^ (Fig. 2a-d). The TEs are sufficiently large such that their variability on the shaft of pyramidal cells has been documented at light microscopy level^36,52^. However, individual spine heads within TEs are small and exhibit a diverse range of sizes and shapes, with spines receiving inputs from different boutons often located in close proximity. Precise quantification requires reconstruction of the spines and their inputs at nanoscale resolution.

**Fig. 2:**
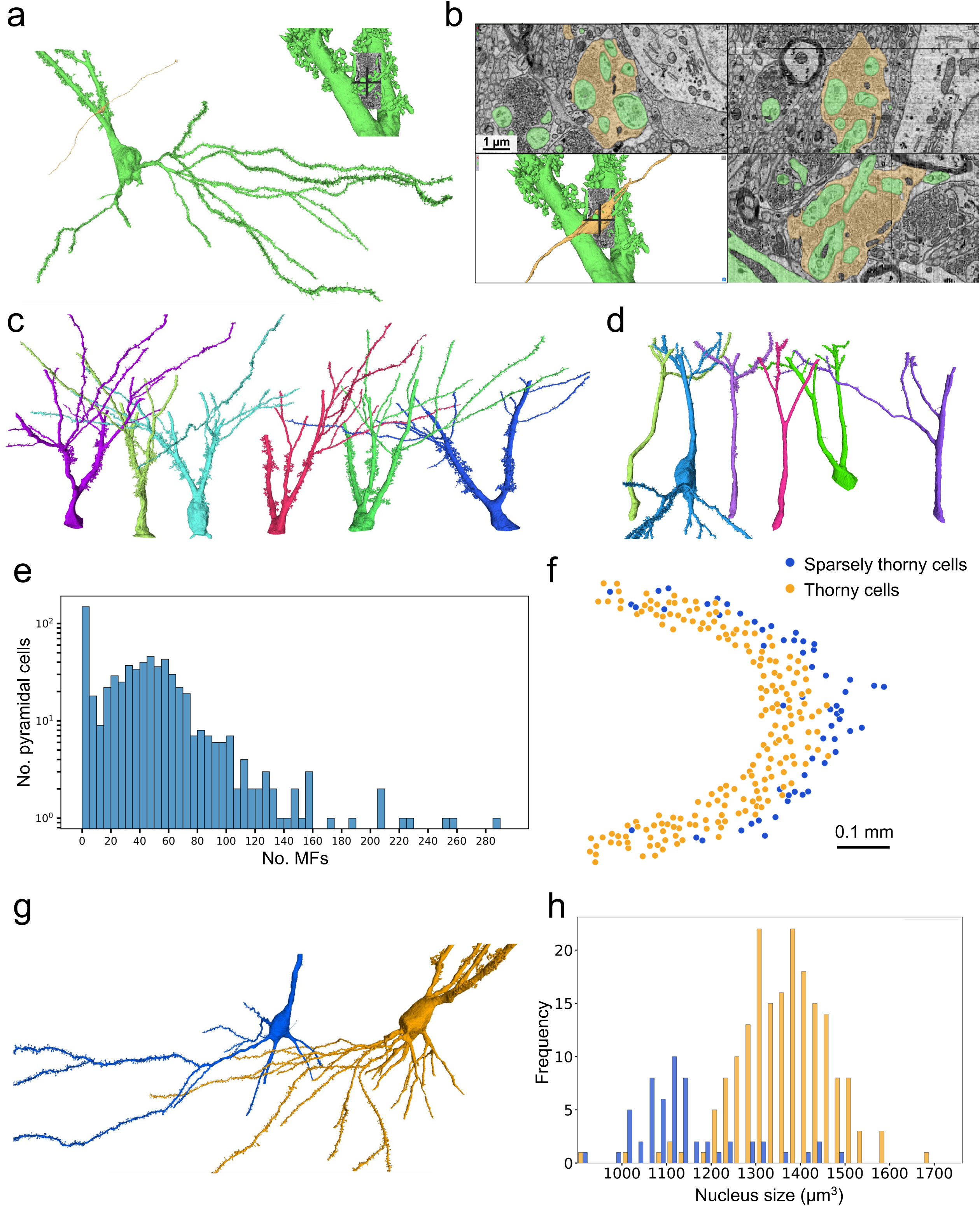
Mapping MF inputs to CA3 pyramidal cells. **a - b,** 3D reconstructions of a MF (yellow) and thorny excrescences of a pyramidal cell (green), with zoom-in views at the xy, xz, yz planes of the CA3 volume. **c - d,** Morphologies of proximal apical dendrites for thorny and sparsely thorny cells. **e,** The number of MFs for long-apical pyramidal cells (nucleus z=1400 - 2143). **f - h,** The spatial distribution of nucleus locations, examples of 3D reconstructions, and a histogram of nucleus sizes for thorny cells (yellow) or sparsely thorny (blue) that are classified based on a threshold of 10 MFs per cell. Only a subset of long-apical pyramidal cells with long apical dendrites and fully contained nuclei within the volume are included (nucleus z=1600 - 1950).

Since the TEs are localized to the proximal apical dendrites close to the soma, we selected pyramidal cells with somata near one end of the Z-axis (z-layers = 1400 - 2143) to maximize coverage of their proximal apical dendrites within the volume. This group is referred to as ‘long-apical pyramidal cells’ (636 cells) and does not imply a distinct cell type. We mapped all MF inputs to the long-apical pyramidal cells, totalling ∼25,200 MFs. While the somata of MFs lie outside the imaged volume, sparse anatomical tracing studies have indicated that each MF axon in CA3 typically originates from a single granule cell^35^. The distribution of the MF inputs is bimodal with a heavy tail (Fig. 2e). One mode comprises a population of CA3 cells (24%) that receive no or minimal MF inputs (sparsely thorny cells). The other mode consists of CA3 cells receiving 15 or more MF inputs per cell, with most having around 50 (thorny cells). This matches with previous indirect estimates for rat^34^. Surprisingly, a small number of CA3 cells (2%) have over 100 MF inputs per cell with the maximum close to 300, many more than previous estimates^34^.

A potential concern is whether the observed distribution is confounded by truncation of the apical dendrites due to the limited z-axis. We observed that the overall distribution and the extreme values of MF inputs are insensitive to the axial location of the nuclei (Extended Data Fig. 4a). Furthermore, a subset of long-apical pyramidal cells (266 cells) have apical dendrites with side branches, which typically mark the distal extent of TEs (Fig. 2c-d), suggesting that most of the TEs are captured. The number of MF inputs to this subset exhibit a similar bimodal distribution (Extended Data Fig. 4b). Therefore, the observed distribution of MF inputs is not significantly biased by volume limitation.

The CA3 pyramidal cells have been classified into two morphologically distinct subtypes, thorny and sparsely thorny, distinguished by abundance of TEs^28–30^. In the CA3 volume, categorizing pyramidal cells solely by their number of MF inputs, using a threshold derived from the distribution (Fig. 2e), recapitulates this previously described division. The cell bodies of the sparsely thorny cells are preferentially located deeper in the pyramidal cell layer (radial axis) and are more at distal region (transverse axis), while thorny cells are situated more superficially within the layer (Fig. 2f). Morphologically, thorny cells have a larger nucleus and more elaborate basal dendrites than sparsely thorny cells (Fig. 2g-h). While our quantification is limited to neurons with both somata and proximal apical dendrites within the volume, selecting cells truncated further away from the soma—thereby capturing more of the upper dendritic arbors—shows that thorny and sparsely thorny cells also differ markedly in the extent of dendritic branching (Extended Data Fig. 4c-e). These morphological distinctions and spatial preferences of two subtypes are in line with previous studies^28,30^.

### MF inputs to CA3 pyramidal cells exhibit a proximodistal gradient

Proximodistal heterogeneity of MF inputs to CA3 has been studied with anatomical tracing and physiological approaches^26,35^. We extended these findings by analyzing how the spatial distribution of pyramidal cells is related to the number of MF inputs they receive. Within the EM volume, the CA3 pyramidal cells exhibit a proximodistal gradient in an increasing number of MF inputs (Fig. 3a). Given pyramidal cells with more MF inputs are localized at the superficial part of the soma layer, the pyramidal cells with most numerous inputs are concentrated at the superficial and distal end (Fig. 3b-c).

**Fig. 3:**
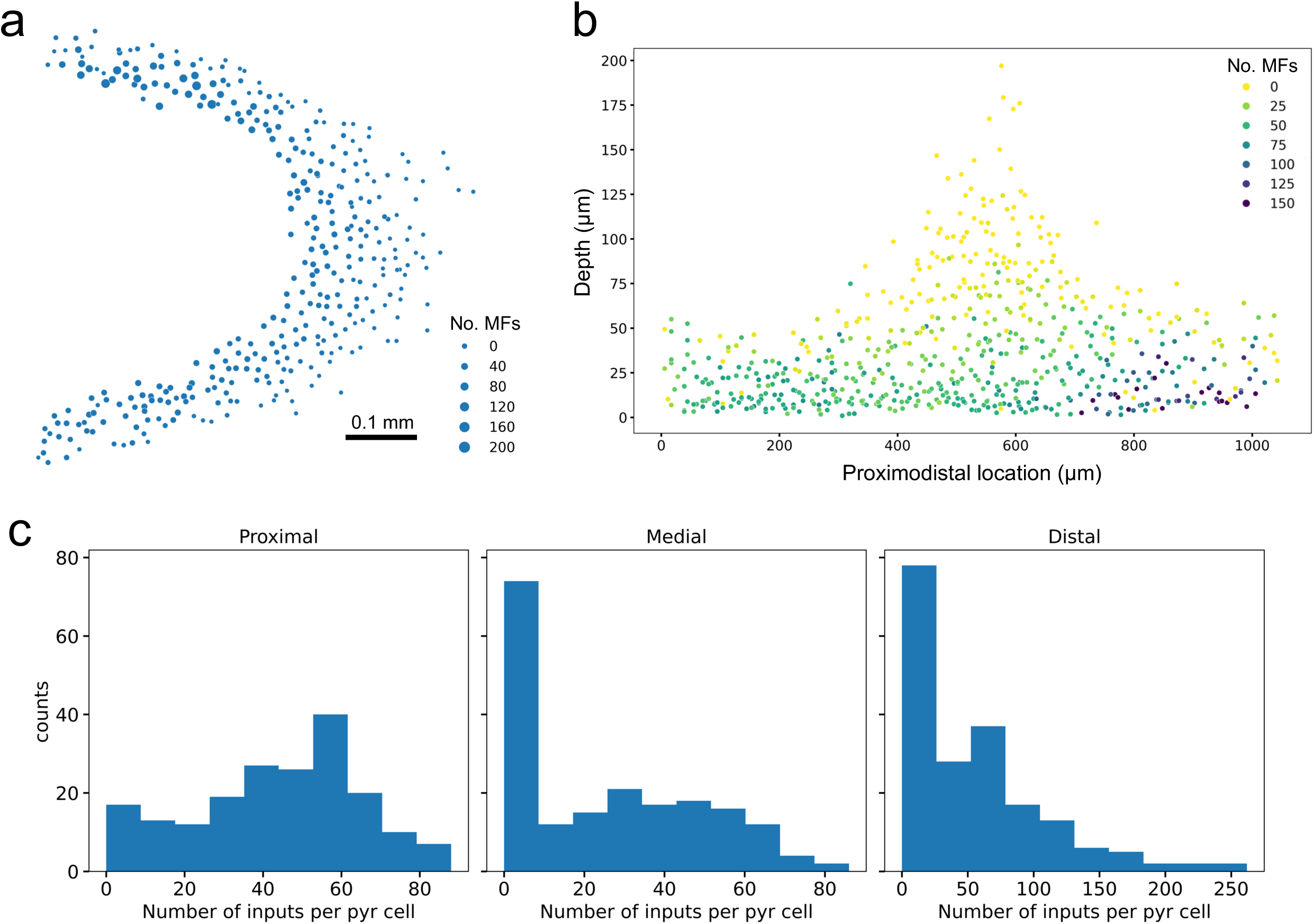
A proximodistal gradient of MF inputs to CA3 pyramidal cells. **a,** Spatial distribution of nucleus locations at xy plane as a function of number of mossy fibers per cell. **b,** Proximodistal and superficial-deep gradients of MF inputs for pyramidal cells. A linear boundary between stratum lucidum and stratum pyramidale is defined such that projected locations of cells on the line indicate proximodistal locations and distance of the cells to the line indicates depth (Methods). **c**, Distribution of the number of MF inputs per pyramidal cell across proximal, medial, and distal regions of CA3. Pyramidal cells were divided into three equally sized groups based on their proximodistal positions.

### Increased sharing of MF inputs in distal pyramidal cells

The MF-Pyr connectivity in CA3 has been hypothesized to participate in pattern separation^17,19,53^. We therefore use theoretical methods developed in pattern separation circuits to analyze their connectivity. Pattern separation circuits are a key stage in the feedforward architecture known as the Marr motif^54,55^, which has analogs in the hippocampus, olfactory system, and cerebellum^18^. Models have been developed to quantify how the connectivity of a two-layered pattern separation circuit determines the dimension of a representation formed by neurons of the second layer^56–58^. High-dimensional representations improve associative learning of the read out neurons, whereas low-dimensional representations facilitate generalization and increase robustness to noise ^57,59^. In pattern separation circuits, dimensionality can be quantified with the participation ratio^56,60^, which measures how uniformly inputs (e.g. MFs) are represented across second-layer neurons (e.g. pyramidal cells) based on their connectivity.

Two key determinants of this dimensionality are randomness and degree—the number of synaptic partners per cell, specifically the number of presynaptic inputs each neuron in the second layer receives. While demonstrating true randomness is inherently challenging, the extent of randomness in a circuit can be quantified through comparisons with defined null models. Applying this framework, analyses of EM-reconstructed connectivity in fly olfactory systems^60,61^ and cerebellum^62^ have found varying levels of deviations from both randomness and uniform degree distributions.

To analyze MF–pyramidal connectivity, we compared the reconstructed network to various null models. While previous models often assume a uniform connection probability^63,64^, connectivity patterns in biological circuits are heavily influenced by additional factors^65^, such as degree distributions (i.e., the number of synaptic partners per cell) and the spatial proximity of synaptic partners. To account for these influences, we compared the frequency of shared MF inputs between pyramidal cell pairs across the observed data, a degree-preserving configuration model^65^, and an anatomically constrained model. The MF-Pyr connectivity can be defined as a bipartite graph, where MF out-degree is the number of target pyramidal cells, and the pyramidal in-degree is the number of presynaptic MFs. The reconstructed connectivity showed more sharing of MF inputs than a configuration model preserving both in- and out-degrees (Fig. 4a-b).

**Fig. 4:**
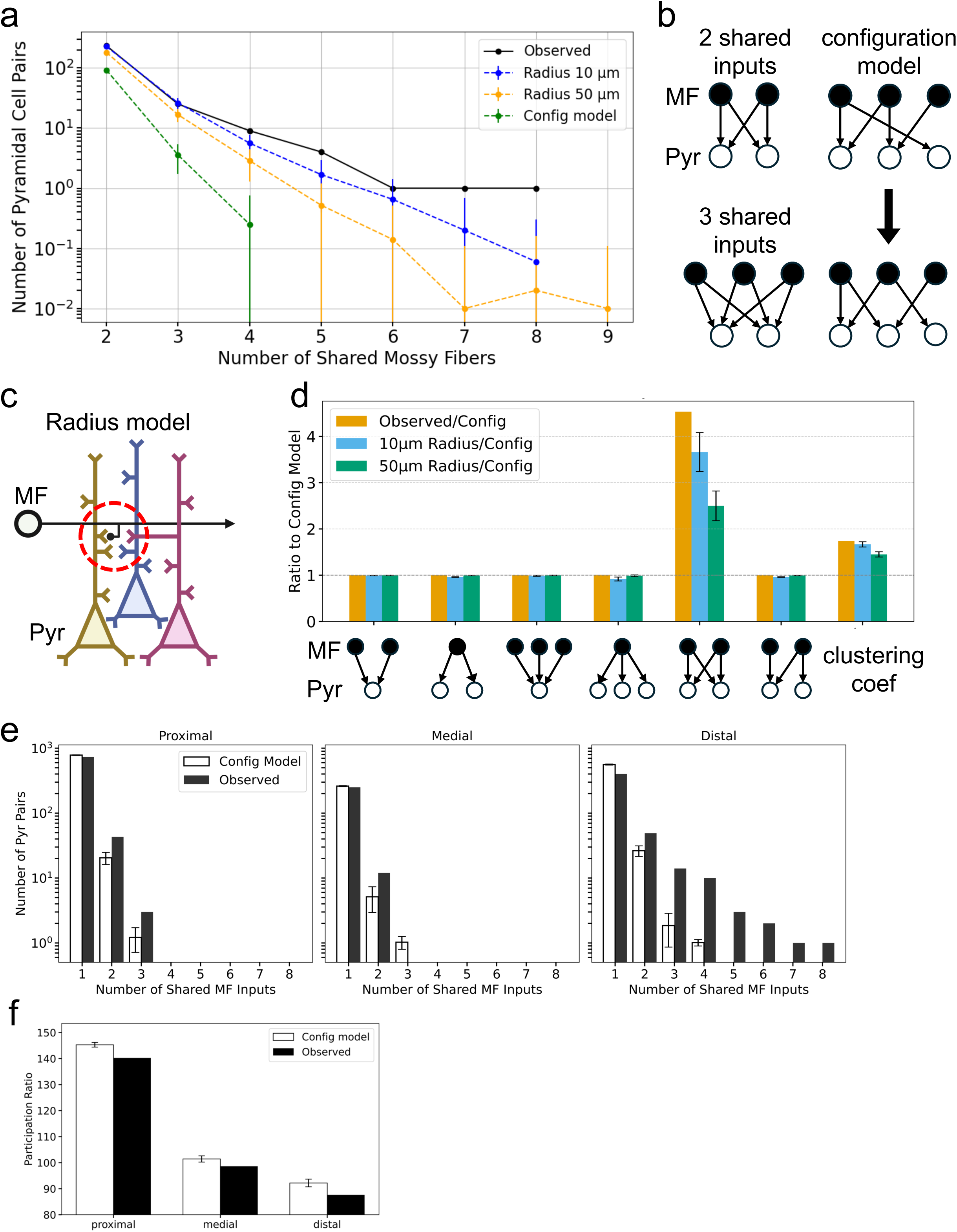
Increased sharing of MF inputs in distal pyramidal cells. **a**, Among 636 long-apical pyramidal cells, the number of cell pairs sharing two or more common mossy fiber (MF) inputs was compared across the observed network, the radius-based models (10 µm and 50 µm), and the configuration model. **b**, Examples of pyramidal cell pairs sharing 2 or 3 mossy fiber (MF) inputs. The configuration model preserves both in- and out-degrees. **c**, In the radius-based model, each MF bouton was randomly connected to thorny spines of pyramidal cells, based on the spatial proximity between the bouton and spines, constrained within a 10 µm or 50 µm radius. **d**, Frequencies of 3-cell and 4-cell motifs, and clustering coefficient (Methods), in the observed network, and in 10 µm- and 50 µm-radius models, shown relative to the configuration model. Error bar denotes 95% confidence interval. **e-f**, For each of the proximal, medial, and distal regions, the number of pyramidal cell pairs sharing two or more common MF inputs (**e**) and participation ratios (**f**; Methods) were compared between the observed data and a configuration model.

Increased common MF inputs could either be clustered onto a subset of pyramidal cells or dispersed across the population. High clustering means if Pyr-a shared inputs with Pyr-b, and Pyr-b shared inputs with Pyr-c, then Pyr-a and Pyr-c are more likely to share inputs. The reconstructed connectivity showed a higher clustering coefficient than a configuration model (Fig. 4d).

Both the increase and clustering of common MF inputs are reduced when the observed connectivity is compared to an anatomically constrained model, in which connections were randomized within smaller radii (Fig. 4a-d; Extended Data Fig. 5a; Methods). Moreover, the distribution of distances between postsynaptic sites of pyramidal cell pairs sharing MF inputs showed a peak at short distances (Extended Data Fig. 5b-c). Indeed, many shared inputs originate from thorny spines of two neighboring pyramidal cells innervated by either the same MF terminal (Extended Data Fig. 5d) or nearby satellite terminals belonging to the same MF (Extended Data Fig. 5e). The spatial gradient observed for MF inputs led us to investigate whether input sharing shows a spatial bias. Increased input was found to occur predominantly among distal pyramidal cells (Fig. 4e).

In addition to forming synapses on thorny spines, the MF terminals also form puncta adherentias (PAs)^1,5^ or synaptic contacts (Extended Data Fig. 5f) on dendritic shafts of pyramidal cells. In the analysis, all connections of pyramidal cell pairs sharing two or more common MF inputs were verified to only include MF inputs onto thorny spines.

In summary, pairs of distal pyramidal cells share more MF inputs than expected from globally randomized connectivity with preserved degrees, but the increased sharing is largely consistent with locally randomized connectivity based on axon-dendrite proximity.

The spatial gradient in MF input connectivity suggests a corresponding gradient in the expected dimensionality of activity in the postsynaptic neural populations. We used the participation ratio^56,60^ of the MF to pyramidal cell synaptic connection matrix to quantify the distribution of neural representation. We observed a decreasing gradient of participation ratios along the proximodistal axis (Fig. 4f). This dimensionality reduction could result from more numerous inputs (i.e. pyramidal in-degrees) or increased input sharing. Comparison with a configuration model that preserves degrees while eliminating input sharing shows only minor differences from the reconstructed data (Fig. 4f). Therefore, increased in-degrees among distal pyramidal cells, rather than input sharing, drives the dimensionality reduction.

### High-convergent pyramidal cells receive inputs via relatively small MF terminals

Dense segmentation at nanoscale allows us to examine not only the number but the strengths of each input, inferred from sizes of synaptic structures. The MF boutons are known to have a variety of sizes and morphologies^45,66^, with anywhere from zero to multiple filopodia extensions. We developed a method of segmenting MF boutons based on the number of neighboring vertices in 3D meshes (Methods; Extended Data Fig. 6a-c), and validated its ability to segment a variety of bouton types, including en passant, terminal, doublets, and boutons with filopodia (Extended Data Fig. 6d). This method was used to segment a majority of MF boutons (24264, > 85% of total) that are connected to the 636 long-apical pyramidal cells. The distribution of bouton sizes has a wide range from a few to over 40 μm^3^ (Fig. 5a), prompting us to examine their relationships with the number of synaptic vesicles. We adapted a deep learning-based cellular segmentation method ^67^ to automatically identify vesicles (Methods; Extended Data Fig. 6e), given the similarity between vesicles and cell bodies from other imaging modalities. The neural network, trained on human-annotated vesicles, achieved high accuracy in predicting vesicles from 2D images of the CA3 volume. Segmentation of vesicles showed that the sizes of boutons have an excellent correlation with the number of vesicles (Fig. 5b), yielding a density of vesicles per volume consistent with previously reported data^5^.

**Fig. 5:**
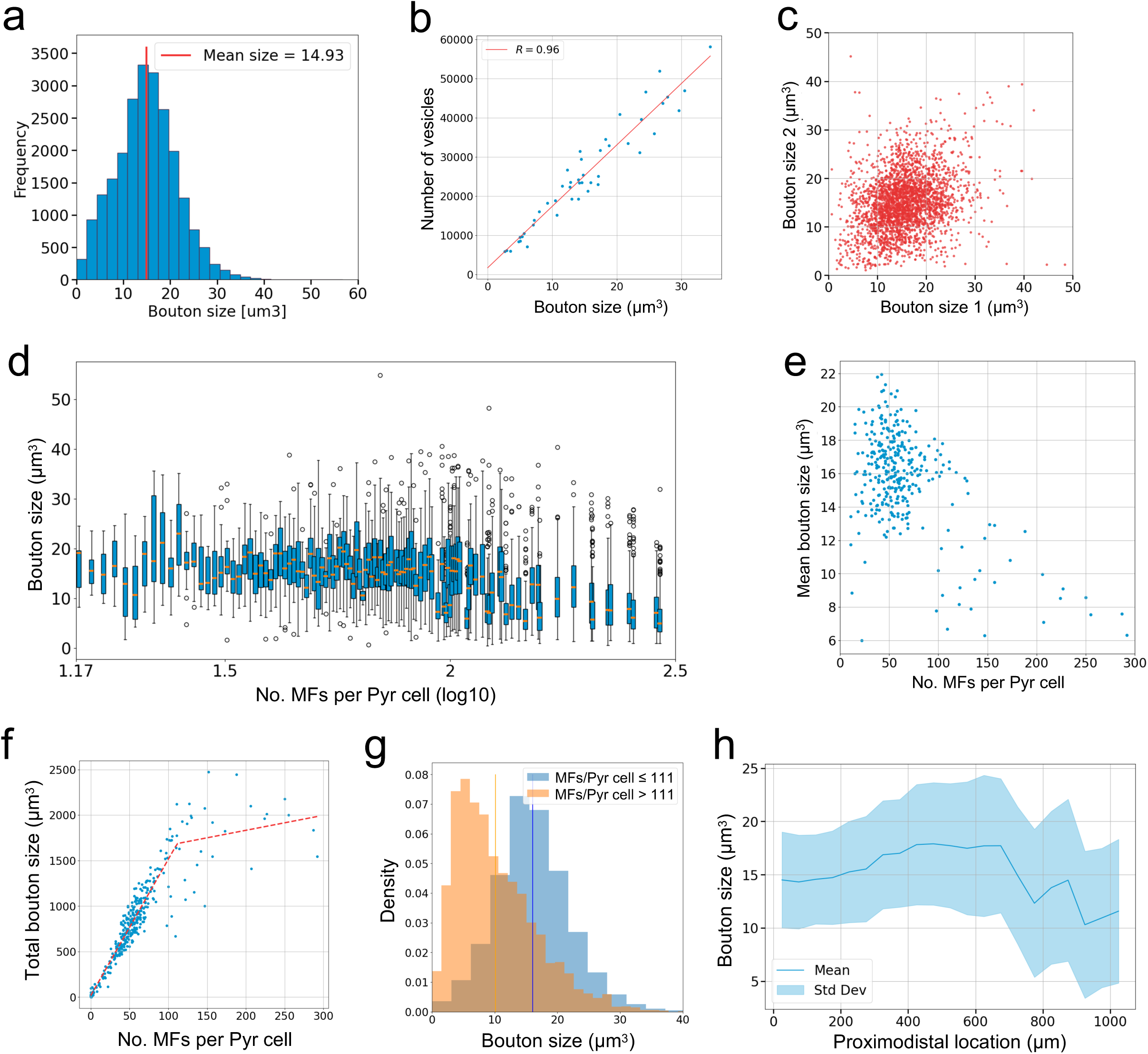
Pyramidal cells with numerous MF inputs are innervated by smaller boutons. **a,** Size distribution of 24,264 MF boutons that are presynaptic to 636 long-apical pyramidal cells (> 85% of their boutons). **b,** Correlation between number of segmented vesicles and boutons size. **c,** Each point represents sizes of two boutons that are from the same MF. **d,** Each box shows the interquartile range (IQR) from the 25th (Q1) to the 75th percentile (Q3), with the median marked by an orange line. Whiskers extend to 1.5×IQR from Q1 and Q3; points beyond are shown as circles. Among pyramidal cells receiving the same number of MF inputs, one was randomly selected for display. Pyramidal cells with less than 15 MF inputs are not shown (log15=1.17). **e,** Mean bouton sizes per pyramidal cell as a function of the number of MF inputs. Data points are shown for pyramidal cells with more than 80% of their boutons segmented. **f,** Summed bouton sizes per pyramidal cell as a function of MF inputs. The data were fitted with a bilinear model, and the inflection point (111 MF inputs) was used as a threshold to separate pyramidal cells into two groups. **g,** Probability density distribution of bouton sizes between high-convergent (> 111 MF inputs) and other pyramidal cells. **f,** Means and standard deviations of MF bouton sizes across proximodistal locations of pyramidal cells, binned in 50 µm intervals.

Analysis of bouton sizes shows that those originating from the same MF are uncorrelated (Fig. 5c). Due to limited volume of the dataset, each MF typically has only 1 to 3 boutons (Extended Data Fig. 6f), while estimates from studies in rats suggest that each MF, on average, forms 11 to 18 large terminals that innervate pyramidal cells^37^. At the postsynaptic side, individual pyramidal cells also receive inputs from boutons with a wide range of sizes (Fig. 5d). Paradoxically, pyramidal cells with high convergence—those with over 111 MF inputs—form synapses with MF boutons that are, on average, smaller in size (Fig. 5e-g). The inverse relationship between MF input number and bouton size may reflect compensatory scaling of synaptic weights. while highly convergent pyramidal cells cluster at the distal end along the transverse axis (Fig. 3b), the average bouton size is smaller distally along the transverse axis (Fig. 5h), matching the reported gradient of decreasing synaptic strength in response to MF stimulation^26^.

### Cell-type-specific feedforward inhibition to pyramidal cells

The classification of pyramidal subtypes allows us to analyze cell type-specific connectivity. To search for specificity in inhibitory connection, we identified segments with the highest number of output synapses in the volume, which are predominantly inhibitory axons originating from neurons with somata outside the imaged volume. We refer to this group as ‘inhibitory axons’ (Extended data Fig. 7a). From this set, we further selected axons with 200 or more output synapses onto pyramidal cells with known subtypes. We found that many of these axons preferentially target either sparsely thorny cells or thorny cells, as indicated by the differing number of synapses onto each group (Fig. 6a). As thorny cells have larger nuclei than sparsely thorny cells (Fig. 2h), when the postsynaptic pyramidal cells were grouped by nucleus sizes, a preferential targeting effect in the number of synapses was also observed (Extended Data Fig. 7b). While this result is consistent with subtype-specific correlated activity observed in physiological recording^29^, the source of inhibition was unknown previously.

**Fig. 6:**
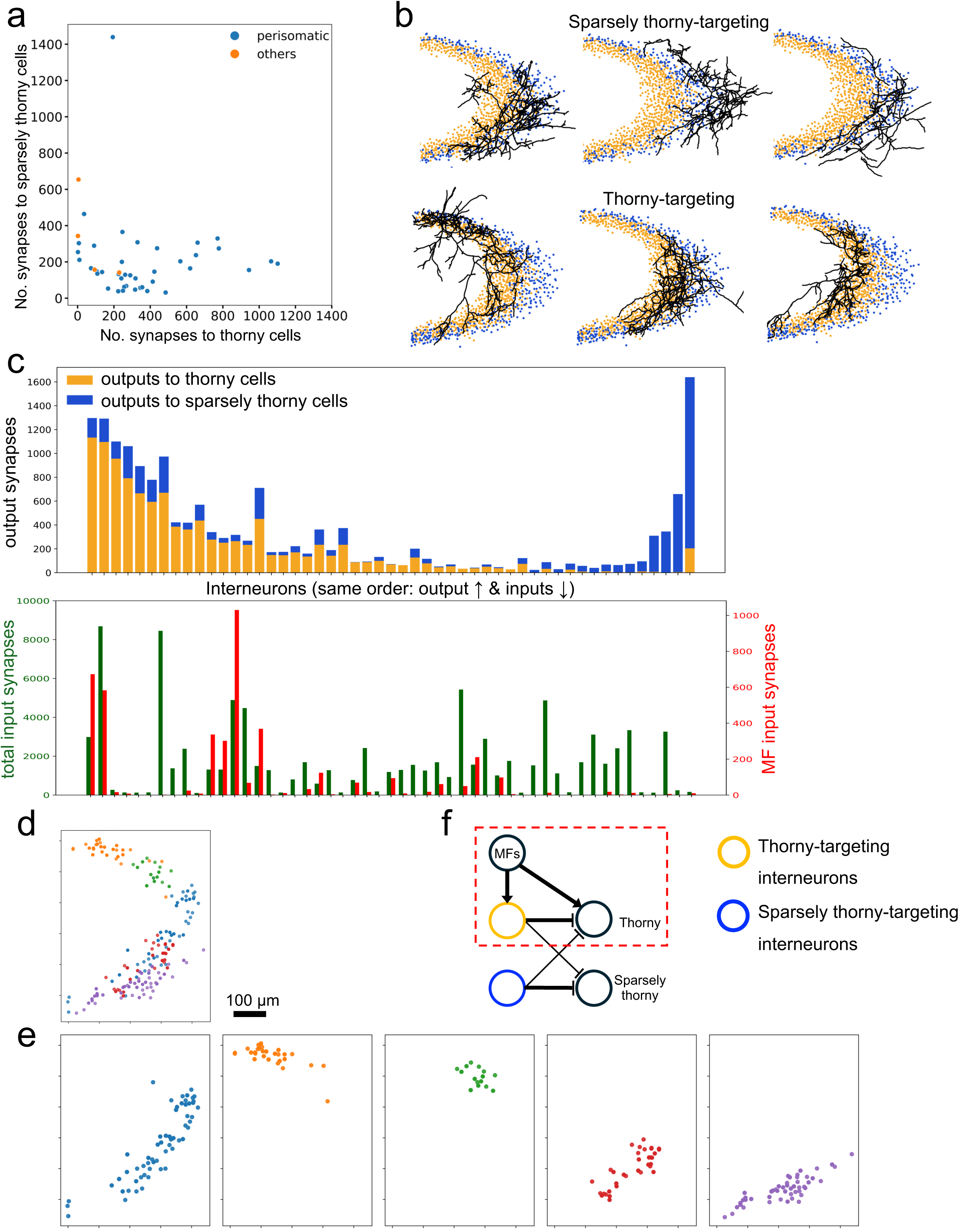
Perisomatic interneurons selectively target thorny and sparsely thorny cells. **a,** Number of inhibitory synapses to thorny (x-axis) and sparsely thorny cells (y-axis). Each circle represents the number of synapses from an inhibitory axon. Inhibitory axons are classified as perisomatic- or dendritic-targeting, based on along-the-skeleton distances from synapses to the nucleus of the postsynaptic pyramidal cells (Methods and Extended Data Fig.6c-d). **b,** Skeletons of inhibitory axons (black) are shown alongside pyramidal cells with large (orange, sparsely thorny cells) and small (blue, thorny cells) nuclei (threshold, 1164 µm^3^). **c**, Only interneurons targeting thorny cells receive MF inputs. Upper panel: number of output synapses from each interneuron onto thorny versus sparsely thorny pyramidal cells. Lower panel: for the same set of interneurons, total input synapse count (left y-axis) and number of MF input synapses (right y-axis) are shown. Output and input data are aligned for each interneuron. **d-e**, Spatial distribution (X and Y coordinates) of pyramidal cell somata involved in feedforward inhibition motifs (red highlighted in **f**), colored by the participating interneuron in each motif. Five interneurons receiving the highest number of mossy fiber inputs are shown together in (**d**) but separately in (**e**). **f**, Diagram of feedforward inhibition motif that selectively targets a pyramidal subtype, thorny cells. Separate interneuron, without MF inputs, targets sparsely thorny cells.

**Fig. 7:**
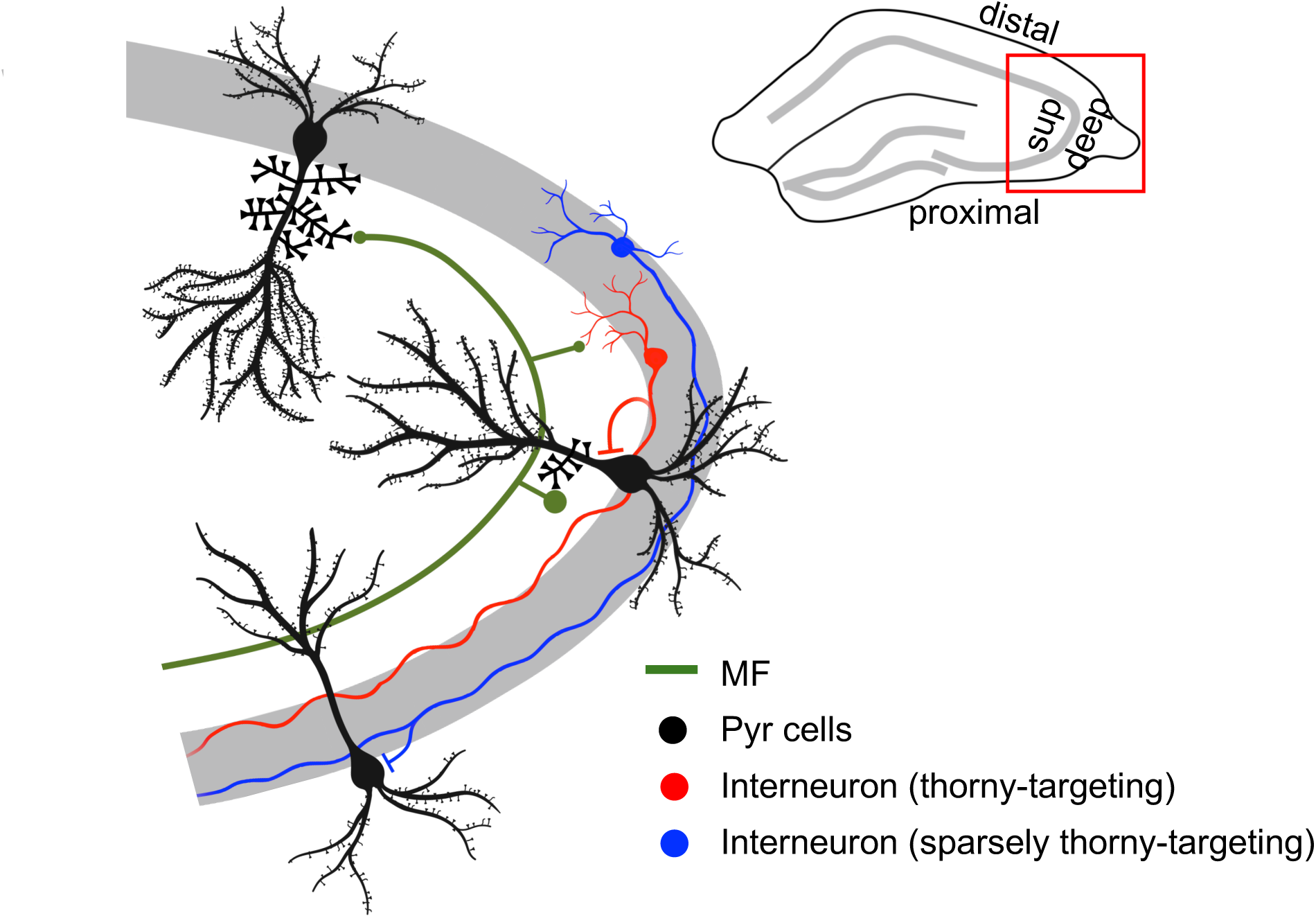
MF-Pyramidal cell connectivity is characterized by high convergence, a spatial gradient, and subtype-specific feedforward inhibition. A schematic highlights our key findings: (1) highly convergent pyramidal cells receive smaller MF terminals; (2) number of MF input follows a proximodistal gradient; (3) a feedforward inhibition circuit including a sublayer-specific pyramidal cell subtype—thorny cells.

A major division of inhibitory cell types is based on their subcellular targeting region: perisomatic versus dendrite-targeting. We measured the along-the-skeleton distances from the postsynaptic sites to the center of the pyramidal cell nucleus. We then analyzed the distribution of synapse-to-soma distances for each inhibitory axon, and determined the mode for each distribution. The overall distribution of modes for all inhibitory axons is bimodal, corresponding to perisomatic and dendrite-targeting groups (Extended Data Fig. 7c-d). A majority of subtype-specific inhibitory axons are found to be perisomatic (Fig. 6a). Notably, the inhibitory synapses from perisomatic inhibitory axons are distributed across many pyramidal cells (Extended Data Fig. 7e-f). The main axonal shafts of many perisomatic inhibitory neurons are myelinated (Extended Data Fig. 8).

Thorny and sparsely thorny cells occupy superficial and deep sublayers of the soma layer, respectively. The preferential targeting of either subtypes by inhibitory axons could arise either from axons projecting selectively to superficial or deep sublayers, or from broadly distributed axonal innervation across the soma layer combined with subtype-specific synapse formation. Examination of axonal projection territory shows that axons of inhibitory cells that preferentially target sparsely thorny cells are predominantly localized to the deeper portion of the pyramidal cell layer, whereas those targeting thorny cells preferentially innervate the superficial portion (Fig. 6b). MFs have been shown to provide abundant outputs to interneurons^37^, which tightly regulate CA3 pyramidal cells through feedforward inhibition^68,69^. We find that MF innervation is selective for interneurons targeting thorny pyramidal cells (Fig. 6c), showing a subtype-specific feedforward inhibition circuit. The pyramidal cells that participated in 3-cell feedforward motifs are regionalized based on the innervation area of interneurons (Fig. 6d-f). In summary, we found a MF-Interneuron-Pyramidal feedforward inhibition motif which is selective for thorny pyramidal cells (Fig. 7).

## Discussion

### Electron microscopy imaging and reconstruction

Acquisitions of connectomic datasets are often dichotomized into SEM- and TEM-based approaches. Automation of sectioning and imaging^9,10,70–72^ has increased the scale of the EM volume for both approaches. However, the largest EM datasets, each at 1 mm³ in volume—one from SEM and one from TEM—still have artifacts^8,12^ which limit the automation and speed of reconstruction. While our current volume (0.1 mm^3^) is modest in size in comparison, we demonstrated here that, with improvements in image quality and computation methods^42,43^, reconstructions from a TEM volume can be obtained with high efficiency and quality. We continue to increase volume size and reduce section loss in our GridTape-TEM acquisition pipeline^41^. Meanwhile, serial-section SEM volumes of comparable quality and larger in size than our volume have recently been reported^11,14,15^, with ongoing reconstruction. The neuroscience community will benefit from healthy competition between the two approaches and, more importantly, a continuing commitment to openness. Building on the success of community-driven connectomics platforms like EyeWire^73^ and FlyWire^74^, Pyr launches as a similarly open platform (Extended Data Fig. 9) dedicated to enable collaborative discoveries of hippocampal connectomics.

### Comparison of CA3 datasets

A recently published 3D EM volume of CA3^14^ is comparable in size to our dataset but differs in several aspects. First, we used a 4-month-old female mouse compared to their 1-month-old male. Second, imaging and reconstruction methods differ. Our dataset is acquired with GridTape TEM while the comparison study used a multi-beam SEM. Our dataset contains slightly more total sections but suffers from more section loss. Notably, we used a deep learning-based alignment and segmentation capable of handling data loss and artifacts and delivering dense segmentation in large-scale datasets^43^. Third, targeted subregions differ (CA3b-c vs. CA3a-b). Proximal CA3 pyramidal cells (e.g. CA3a), targeted by the previous study^14^, are thought to be smaller and contain less recurrent collaterals than distal CA3 cells (e.g. CA3b,c)^75,76^ that are contained in our study. Fourth, the sectioning planes differ. Sammons et al. sectioned more parallel to the apical dendrites, capturing fewer somata but spanning all CA3 layers. In contrast, our dataset is angled closer to orthogonal to the apical dendrites, missing parts of stratum radiatum and most of the molecular layer but including more cell bodies in the transverse plane, allowing for analysis of systematic variations along this dimension. Plurality in animal characteristics, imaging modalities, reconstruction methods, and anatomic areas provide complementary perspectives for connectomic studies.

### Systematic variation of neuronal properties in hippocampus CA3 and CA1

While the heterogeneity of hippocampal pyramidal cells has been well recognized^77^, growing evidence suggests that many neuronal properties vary in a systematic manner. One notable example is the superficial–deep axis in CA3, which exhibits consistent division in dendritic morphology^30^, electrophysiological properties^28,78^, genetic markers^79^, birth timing^80^, and physiologically assessed connectivity^29,31^. Our study extends this organizational principle to connectivity at the ultrastructural level, by quantifying the number of MF inputs and revealing sublayer-specific inhibitory inputs. Furthermore, along the proximodistal axis, we identified a gradient in both the number and size of MF inputs, adding to known variations in morphology^23^, physiology^26^, and functions^24,25^.

In the CA1 region, superficial versus deep pyramidal cells have been studied for differences in physiology^81^, long-range entorhinal inputs^82^, transcriptional profiles^83^, birthdating^84^, spatial activities^85,86^, and inhibitory input^87^. Together, these findings point to a common principle of layered organization across hippocampal subfields.

### Functional implication

Sparsely thorny and thorny cells exhibit regular spiking and bursting activity, respectively, and the bursting propensity of sparsely thorny cells was essential for initiating sharp waves events—synchronized activity patterns important for memory function in the hippocampus^28^. Our findings show that perisomatic inhibitory axons target pyramidal subtypes with highly distributed synapses (Extended Data Fig. 7e-f), suggesting inhibitory cells synchronize the activity of a group of cells within each subtype. Furthermore, the inhibitory axons are myelinated, suggesting a role for fast, temporally precise inhibition in coordinating network activity during sharp wave events ^88–90^.

Studies of spatial firing properties along the CA3 transverse axis have suggested a functional gradient, with strong pattern separation in proximal CA3 and robust pattern completion in distal CA3 ^24,25,27^. The underlying circuit mechanism was thought to be a graded recurrent connectivity in the proximodistal direction, as recurrent collaterals are more extensive distally^26,75,76^. Analysis of recurrent connectivity requires circuit analysis and dataset-wide segmentation of synaptic weights (e.g., spine head volumes), while accounting for the limitation posed by neuron truncation—particularly of axons. This remains a topic of interest for follow-up study.

We found another proximodistal gradient in the number of MF inputs per pyramidal cell, with a few distal cells receiving well over 200 MF inputs. Greater MF convergence may enable distal CA3 to integrate a broader range of input patterns. This finding is consistent with the larger place field sizes reported in distal compared to proximal CA3^24,25^. Using previously developed metrics^56,60^, we showed that more numerous inputs in pyramidal cells corresponds to a reduction in dimensionality of representation along the same proximodistal axis (Fig. 4e-f). Further modeling studies are needed to explore the relationship between high convergence and the proposed functional roles of CA3 such as pattern completion. Additionally, directly relating functional properties of CA3 cells to number of inputs will require combining *in vivo* imaging with connectomic mapping on the same cells.

### Larger electron microscopy volume

Together the recent datasets^14,16^ represent the largest hippocampal EM volumes currently known. This scale, however, pales by comparison with the expansive, complex axons of CA3 pyramidal cells. In rodents, the hippocampus forms a long, curved structure spanning the brain’s dorsal-to-ventral extent—the longitudinal axis. Reconstructions of single neurons at brain scale have shown that CA3 axons not only span much of the longitudinal axis but also project to and extensively innervate the contralateral hippocampus in both CA3 and CA1 regions^91^. The full extent of these projections far exceeds the scale of current largest EM volumes at 1 mm^3^. The volume size limitation is particularly pronounced along the Z-axis, due to the limited number of serial sections. A significant scale-up of imaging technology over an order of magnitude in size up to a whole mouse brain^92^, is needed to reconstruct the connectivity of these brain-spanning neurons in full.

## Methods

### Sample preparation

#### Animal

All procedures were carried out in accordance with the Institutional Animal Care and Use Committee at Princeton University. A subset of un-aligned sections from this sample have been used previously^41^. A female mouse (C57BL/6J-Tg(Thy1-GCaMP6f) GP5.3Dkim/J, Jackson Laboratories, 028280) aged 4 months was transcardially perfused with a 4 °C fixative mixture of 2.5% paraformaldehyde and 1.3% glutaraldehyde in 0.1M Cacodylate with 2mM CaCl_2_ pH 7.4. The brain was extracted and post-fixed for 36 hrs at 4 °C in the same fixative solution. The perfused brain was subsequently rinsed in 0.1M Cacodylate with 2mM CaCl_2_ for 1 hr (3 x 20 mins, 4 °C) and 300 μm coronal sections were cut on a Leica Vibratome on ice.

#### EM staining

Sample blocks at 300 μm thick were cut out and stained based on a modified reduced osmium (rOTO) protocol with the addition of formamide^40^. The tissue blocks were first *en bloc* stained with 8% formamide, 2% osmium tetroxide, 1.5% potassium ferrocyanide for 3 hours at room temperature. Subsequently, the samples were immersed in 1% thiocarbohydrazide (TCH, 94 mM) 50 °C for 50 mins, followed by a second step of 2% osmium staining for 3 hours at room temperature. The sample was placed in 1% uranyl acetate overnight at 4 °C, followed by lead aspartate (Walton’s, 20 mM lead nitrate in 30 mM aspartate buffer, pH 5.5) at 50 °C for 2 hours. After being washed with water (3 x 10 mins, room temperature), samples proceeded through a graded acetonitrile dehydration series (50%, 75%, 90% w/v in acetonitrile, 10 minutes each at 4 °C, then 4 x 10 minutes of 100% acetonitrile at room temperature). After a progressive resin infiltration series (33% resin:acetonitrile, 50% resin:acetonitrile, 66% resin:acetonitrile, 8 hours each at room temperature), the sample was incubated in fresh 100% resin overnight at room temperature and the resin was cured in the oven at 60 °C for at least 48 hrs. Embedded tissue blocks were then imaged by x-ray micro-computed tomography (Zeiss Xradia Versa 520) to evaluate staining quality and tissue integrity. The sample was imaged with an 4x optical magnification, yielding a pixel size of 1 - 2 µm. The tissue block that best contains the target brain region (hippocampus CA3) was chosen for serial sectioning.

#### Sectioning

The tissue block spans several millimeters in the XY plane, encompassing CA1, CA2, CA3, and part of DG. The main distinction between CA2 and CA3 is the stratum lucidum—a cell-free layer that is difficult to discern using light microscopy. X-ray μCT imaging of EM stained tissue blocks can resolve individual nuclei, fine blood vessels, and in this case, a slightly more translucent stratum lucidum layer immediately superficial of the soma layer. Additionally, the resolution allows identification of the CA3 region by its comparatively sparse nuclei and unambiguous correspondence to the same structures imaged in EM (Fig. 1a-b). We therefore first imaged the block with μCT to precisely locate the CA3 region for subsequent serial sectioning.

The target sample block was first trimmed to a mesa containing the CA3 region using an ultramicrotome (Leica UC7) and a diamond trimming knife (Trim 90, EMS-Diatome). The mesa was trimmed into an oblong hexagonal shape of 4.5 mm height and 1 mm width such that the section fully covers the aperture sized 2 mm x 1.5 mm (GridTape, Luxel) and contains the target tissue at the center of the section. To facilitate adherence of the sections to the aperture on the tape during collection, the leading edge tip was trimmed to have an obtuse angle and the trailing edge tip was trimmed to have an acute angle. We combined an automated tape collecting system (ATUMtome, RMC/Boeckeler) and an ultramicrotome (UC7, Leica) to create a sectioning and collection system and further modified it to be compatible with GridTape ^41^. The series was cut with a 4 mm wide 35 degree ultra Maxi Diamond Knife (Diatome). The cutting speed was set to 0.3 mm/s at a nominal thickness of 40 - 45 nm, and the speed of the tape was synchronized with this cutting rate through a closed-loop system. A total of 2145 sections were cut. The typical cycle time of cutting and collecting one section is 20 seconds, resulting in a total sectioning time of about 12 hours for the entire series.

### TEM imaging

The sections were imaged using two JEOL-1200EXII 120-kV TEMs customized with a beam deflection mechanism (Voxa), a reel-to-reel tape translation system (Voxa), and a lens assembly (AMT, NanoSprint50M-AV), and a high speed CMOS camera (XIMEA, CB500MG-CM). At each stage location, the beam deflection was configured to image 9 sub-tiles, each sized at 6000 x 6000 pixels with a pixel resolution of ∼2.9 nm. Each image tile contains 36 megapixels (or megabytes) of data. Each set of 9 tiles forms a supertile, with approximately 900 pixels of overlap between neighboring tiles within each supertile—equivalent to about 15% of the tile dimensions. The nominal overlap between sub-tiles of neighboring supertiles is approximately 600 pixels—equivalent to about 10% of the tile dimensions. The imaging area per section is around 1 mm x 1 mm to be in excess of the whole tissue section. A section typically comprises 440 supertiles of 20 by 22. With 9 tiles per supertile, this yields a typical total of 3960 tiles for a section. The whole dataset has over 8.5 million tile images. Imaging of the series took place over a period of about 5 months as the imaging software and hardware was being developed. The typical size of raw data for each section is ∼133 GB, totaling over 280 TB of data for the whole dataset.

From the 2145 sections, the beginning part (Z = 0 - 95) was seriously damaged and therefore not used for the segmentation. The main series, which was aligned and segmented, spans 2048 layers in the z-axis (Z = 96 to 2143, inclusive). Within this core series, consecutive losses include one instance of five-section loss, one instance of three-section loss, three instances of two-section loss, and 25 instances of single-section loss (Extended Data Fig.1a). Most of the consecutive section loss occurred during the imaging process. The primary causes were precipitous venting of the microscope column and incomplete drying of residual water on the sections. We have since performed maintenance and modified hardware to address these issues. After alignment and segmentation, the resulting size of the volume is 1 mm x 1 mm x 92.2 µm.

While our segmentation quality and proofreading speed are competitive in large-scale datasets, most neurons are truncated, particularly along the z-axis (Extended Data Fig. 3d). We used morphological features—such as apical side branches—to ensure complete mapping of MF inputs, and separately examined the upper apical dendrites of thorny and sparsely thorny cells whose lower apical dendrites were truncated (Fig. 2c-d). The dominant cause of segmentation errors is missing sections (Extended Data Fig. 2d), especially across a five-section gap. However, most processes can be followed across the five-section gap, although proofreading is required (Extended Data Fig. 1b-c). Moreover, circuit reconstructions in other mammalian TEM volumes with a comparable number of consecutively missing sections have also been reported^62,93^. Efforts to improve both the volume scale and lost-section rate in our EM pipeline are currently underway.

### Montaging

The processing of assembling the volume includes stitching and alignment. Stitching (i.e. montaging) refers to montaging of all image tiles within a section via the overlap between neighboring tiles. Stitching was done using a package based on an elastic spring mesh algorithm (AlignTK, https://mmbios.pitt.edu/aligntk-home). In summary, adjacent images are first registered through a linear transformation followed by a non-linear mapping based on image correlation. Each image is represented as a grid of nodes, with pairwise mappings represented by springs connecting nodes. Consequently, each section is represented as a large spring network, and a solution is derived through a multi-resolution relaxation algorithm. Stitching computation can generally be parallelized in two ways. First, correlation is computed between pairs of image tiles and can be distributed across multiple processes in a computing node. Second, because stitching operates on a per-section basis, each section can be independently assigned to a separate computing node for parallel processing. Processes within each node are concurrently executed using the parallel processing mechanism (mpirun) in the alignTK package. Before rendering for subsequent alignment, image tiles are contrast normalized using the gen_imaps function. Montaging of each section, when run in parallel with other sections, typically takes 4 - 6 hours and includes pairwise correlation computation, mesh relaxation, contrast normalization, manual inspection, and rendering. Montaging of the whole series takes over 4 months in total.

### Segmentation

Images were aligned into a 3D stack by a vendor (Zetta AI) using published methods^42,94,95^. The vendor also pretrained a residual symmetric U-Net^96,97^,with data augmentation^43^ to predict affinities between neighboring voxels at 16×16×40 nm³ manually annotated data derived from three mouse visual cortex datasets^12^. The U-Net was fine-tuned on manually annotated data from the present dataset at 18×18×45 nm³.

Segmentation used methods described previously^43^. The only modification was the addition of a multi-resolution approach^44^, in order to reduce merge errors between large objects caused by image defects and broken cell membranes. We generated a conservative low-resolution segmentation at 72×72×45 nm³, using a low-resolution affinity model trained on a downsampled 18×18×45 nm³ segmentation of a small cutout. We upsampled the low-resolution segmentation using nearest-neighbors to 18×18×45 nm³ and used it as a constraint for the high-resolution segmentation at 18×18×45 nm³. Specifically, the upsampled low-resolution segments were treated as semantic labels, and agglomerations between fragments with different labels were forbidden. The results may contain additional split errors but no new merge errors between large objects. The final step of the segmentation pipeline treated the latest high-resolution segments as supervoxels and agglomerated them without the low-resolution label constraint. This step fixes large split errors where fragments have different low-resolution labels. Instead, we added semantic constraints for predicted compartment labels and size heuristics to prevent unwanted mergers^43^.

The segmentation bounding box measures 51,421 × 57,385 × 2,048 voxels, with a voxel size of 18 × 18 × 45 nm, corresponding to a physical volume of 0.93 × 1.03 × 0.092 mm, or 0.088 mm³. Due to irregular edges and black space in the upper left and right corners of the volume, the total number of segmented voxels is approximately four trillions.

### Synapses

The vendor trained two residual symmetric U-Nets, one for synaptic cleft detection at 18×18×45 nm³ and one for presynaptic and postsynaptic mask predictions at 32×32×45 nm³ ^98^. The models were trained on manually annotated data sampled from various regions across the present dataset.

The cleft probability map was downsampled by 2×, thresholded, and segmented with connected components. Segments with fewer than 5 voxels were excluded. Presynaptic and postsynaptic cells for each cleft were assigned following^98^.

### Reconstruction Pipeline

The inferences were done using a modified version of Chunkflow^99^, segmentation was created using Abiss^100^, and synaptic clefts and assignments were generated with Synaptor^98^. The reconstruction pipeline was orchestrated by SEURON (https://github.com/seung-lab/seuron). The segmentation and synapses were ingested into the CAVE proofreading platform for further analysis^39^.

### Quantification of segmentation accuracy

#### Segmentation failure modes

For a semi-quantitative description of failure modes, we randomly chose 100 segmentation errors and attempted to attribute them to various causes (Extended Data Fig. 2d). The leading cause was consecutive missing sections. With appropriate training data augmentation, a convolutional net can detect neuronal boundaries with high accuracy in spite of missing sections^96^. But accuracy plummets if several consecutive sections are missing. Other failure modes include inaccuracies of the segmentation model to identify intracellular organelle (Extended Data Fig. 2d-e) or extracellular space (Extended Data Fig. 2f). Misalignment now occurs at a rather low rate and is no longer a leading cause of segmentation errors. However, some segmentation errors are perhaps not caused by a single factor, but rather by a “perfect storm” of a failure mode in Extended Data Fig. 2d, a small misalignment, and a neurite that is almost parallel to the plane of the section.

#### Mean axon path lengths between errors^101^

For each neuron, we computed the axon path length between errors by dividing the total skeleton length of the fully proofread axon by the number of proofreading edits (Fig. 1d). The mean axon path length between errors was 108 µm. As there are cases where multiple edits were required to correct a single error, this number is a lower bound. Only pyramidal cell axons extending from soma were used. The initial portion of axons near the soma tend to be thicker and less prone to segmentation errors. To reduce bias, we restricted analysis to pyramidal cells with axons longer than 0.5 mm.

#### Evaluation of continuity through missing sections

We examined the largest gap in the dataset (five missing sections, Z=1262-1266, inclusive) by exhaustively proofreading nearly 500 segments within two randomly selected regions, one densely populated with axons and the other with dendrites and spines (Extended Data Fig. 1b-c). In all cases, the segments—including fine processes—could either be traced through the gap (> 95%) or terminated. We concluded that our proofread segmentation is highly accurate in spite of the missing sections. Furthermore, it may be possible in the future to automate segmentation across gaps of as large as five sections, at least for data of sufficiently high resolution and contrast.

To assess the continuity of neuronal processes across this gap, we selected two regions of interest (ROIs): one densely populated with axons (Extended Data Fig. 1b), and another containing a mixture of thin axons, spines, and dendrites (Extended Data Fig. 1c). In both ROIs, all neurites were exhaustively proofread in the sections immediately adjacent to the gap (z = 1261 and z = 1267). We identified 322 neurite segments in the first ROI and 162 in the second. For each of all segments, we were able to either trace a likely continuation across the gap or determine that the segment represented a spine head terminating near the gap boundary.

### Proofreading

#### Proofreading infrastructure

The automated segmentation was ingested into the ChunkedGraph data structure, and the Connectome Annotation Versioning Engine (CAVE) is used for hosting the proofreadable segmentation and annotations^39^.

#### Proofreading rate

Proofreading rate was computed by dividing the skeletal length of the final proofread neuron segment over proofreading time. To calculate total proofreading time per neuron, the timestamps of proofreading events are extracted and the time intervals between consecutive proofreading events that are less than or equal to one hour apart are summed for each neuron. Anything over an hour between proofreading events was interpreted as a stop by the proofreader. When calculating the skeleton length, branches shorter than 100 µm were excluded. The proofreading rate is calculated by dividing the total skeleton length by the total proofreading time. Pyramidal axons are thinner than dendrites and more prone to segmentation errors. Therefore, only axon proofreading events and axon lengths were used to assess proofreading rate.

### Neuron classification

#### Excitatory and inhibitory neurons

Neocortical neurons can be distinguished from non-neurons largely by the size of the nucleus. In the current CA3 volume, the automated nucleus segmentation provides a list of all nuclei along with their sizes. The distribution of nucleus sizes shows several peaks that largely account for non-neuronal cells, inhibitory neurons, and excitatory neurons (Fig. 1e). The list of cells from each peak were then used as candidates for further classification based on morphology. The classification of excitatory neurons (i.e. pyramidal cells) and inhibitory neurons is based on specific features of pyramidal spines as well as their density. In general, dendrites of pyramidal cells are densely decorated with spines (Extended Data Fig. 2a) while dendrites of inhibitory cells are smooth. While the hippocampus has been shown to have inhibitory cells that have spines^102^, their spines are typically sparser, thinner, and longer than those found on excitatory neurons (Extended Data Fig. 3b). Inhibitory subtypes were further classified based on the number of output synapses in each layer (Extended Data Fig. 3c).

#### Pyramidal subtypes

Due to limited range along the z axis, different cellular components—such as the apical dendrite, nucleus (soma), and basal dendrite—are truncated to varying degrees, depending on the position of the cells within the dataset (Extended Data Fig. 3d). Therefore, different groups of thorny and sparsely thorny cells are selected for different analyses, depending on which components are fully contained within the volume.

In Fig. 2e, the goal is to enumerate all MF inputs of each pyramidal cell. Therefore, 636 ‘long-apical pyramidal cells’ were selected to maximize the length of proximal apical dendrite and are often truncated at the level of the soma or nuclei, with basal dendrites outside of the volume. The distribution of MF inputs to pyramidal cells is bimodal, with one peak near zero and another centered around 50 (Fig. 2e). A threshold of 10 MF inputs—corresponding to the trough between the two peaks—was used to classify pyramidal cells as either thorny or sparsely thorny. In Fig. 2f-h, the goal is to analyze the relationship between nucleus size and the two subtypes. Therefore, a smaller subset of 240 pyramidal cells were selected that contain both relatively long apical dendrite and complete nuclei. In Fig. 6a, the goal was to analyze perisomatic inputs to pyramidal cells. Because perisomatic, inhibitory inputs are distributed (Extended Data Fig. 7e-f), we tried to maximize the number of cells that have soma in the volume and can be identified as thorny or sparsely thorny (754 pyramidal cells). These cells were classified based on at least two of the following criteria: (1) if more than 10 MF inputs were identified, the cell was classified as thorny; (2) a nucleus size threshold of 1150 µm^3^ with small nuclei corresponding to sparsely thorny cells, and large nuclei to thorny cells; (3) a combination of additional morphological features—known to differ between thorny and sparsely thorny cells^28,30^—was used, including depth in the soma layer, size of the apical dendritic trunk, length of unbranched dendritic shaft, and the complexity of basal dendrites (Extended Data Fig. 4c-e).

### Laminar organization and spatial axes

We defined four layers in the dataset: stratum radiatum (s.r.), stratum lucidum (s.l.), stratum pyramidale (s.p.), and stratum oriens (s.o.) (Fig.1f). The pyramidale (s.p.) was defined by the spatial distribution of pyramidal cell bodies, identified via spatial locations of the centroids of their nuclei. The lucidum (s.l.) was delineated by the presence of large MF–pyramidal (MF–pyr) synapses, identified by thresholding aggregated synapse size at 10,000 voxels, with a voxel size of 18 × 18 × 45 nm³. We modeled the 3D curved layer boundary between s.l. and s.p. layers as a nonlinear decision boundary based on these two point distributions, using the Support Vector Classification (SVC) algorithm from the Python package scikit-learn and a nonlinear classifier with a radial basis function (RBF) kernel. This nonlinear boundary effectively distinguished the spatial distributions of pyramidal cell nuclei (s.p.) and MF-Pyr synapses (s.l.). The deeper region lacking pyramidal cell nuclei was defined as s.o., while the superficial region lacking large MF–pyr synapses was defined as s.r.. We projected this 3D nonlinear boundary onto the XY plane to analyze the locations of the pyramidal cells along the proximo-distal axis extending from the DG side to the CA2 side. A quartic polynomial curve was fitted to the projected 2D boundary, mathematically defined by the following equation: y = 3.922e-13 x⁴ - 7.211e-08 x³ + 0.005003 x² - 155.2 x + 1.869e+06.

To quantify the spatial positions of pyramidal cell bodies, we defined two metrics: curve distance and depth. Curve distance refers to the distance along the fitted quartic polynomial curve, starting from the point of the curve that is nearest to the DG. Curve distance was therefore used as a metric of proximodistal location. Each pyramidal cell nucleus was mapped to its nearest corresponding point on the polynomial curve. Depth was calculated as the Euclidean distance between each nucleus centroid and the closest point on the 3D boundary between s.l. and s.p. Layers.

### Identification of MF-pyramidal cell connectivity

The MF inputs to pyramidal cells were identified based on the following criteria: (1) localization within the stratum lucidum, (2) envelopment of the postsynaptic structure by a large presynaptic bouton, and (3) the presence of two or more clustered thorny spines. During analysis of MF–pyramidal (MF–Pyr) synapses, we observed numerous contacts between MF terminals and the dendritic shafts of neighboring pyramidal cells. We considered some of these shaft contacts to be synaptic, as they displayed characteristic features such as docked synaptic vesicles, asymmetric postsynaptic densities, and, in some cases, an absence of other plausible postsynaptic partners. However, we also noted that some shaft contacts likely represent a form of membrane specialization known as puncta adherentia^1,5^. To ensure that the analyzed connectivity reflects only MF inputs onto thorny spines, we manually inspected all MF–Pyr contacts (∼2,700) that contributed to two or more shared MF inputs (Fig. 4a), and excluded any that did not meet thorny spine-based synaptic criteria.

### Bouton extraction and vesicle segmentation

We addressed the bouton extraction problem using a 3D mesh vertex classification method. First, given a MF, its postsynaptic pyramidal cell, and the coordinates of synapses connecting these two cells, we constructed an initial bounding box around the synapse coordinates. This bounding box was subsequently enlarged by 1000 voxels along the X and Y axes, and 400 voxels along the Z axis with a voxel size of 18 × 18 × 45 nm^3^. Within the enlarged bounding box, we loaded the 3D mesh representation of the MF including the bouton (Extended Data Fig. 6a). Each vertex of the mesh was classified based on local voxel density. For each vertex, we counted the number of neighboring vertices located within a radius of 2.5 μm. Due to the swelling nature of boutons, the distribution of vertex counts exhibited a bimodal distribution (Extended Data Fig. 6b). To determine the decision boundary for classification, we automatically identified a threshold at the minimum point between the two peaks in the bimodal distribution. Vertices with a neighbor count larger than the threshold were classified as bouton vertices (Extended Data Fig. 6c). The volume of each bouton was measured by taking the 3D convex hull of the bouton vertices and counting the number of segmentation voxels contained within the hull. Occasionally a single MF forms multiple boutons targeting the same postsynaptic cell. If the MF has at least two synapses located more than 5 μm away from the centroid of all synapse coordinates, we concluded that the MF likely forms multiple boutons. In such cases, we determined the number of boutons by clustering the synapse coordinates in 3D space using k-means clustering (k=2, 3 and 4). The best number of clusters was selected based on the k-means inertia score. For each synapse cluster, we constructed an initial bounding box around the cluster. If the synapse cluster contained fewer than 24 synapses, the bounding box was enlarged by 400 voxels along the X and Y axes and 160 voxels along Z axis. Otherwise, the box was enlarged by 600 voxels along the X and Y axes and 240 voxels along the Z axis.

To demonstrate the validity of the bouton extraction, we applied it to segment boutons of various cases of the bouton shape, which include en-passant, terminal, T-junction, small, medium, large, filopodial, and double boutons (Extended Data Fig. 6d). The method segmented the bouton correctly in all 100 randomly selected test cases, excluding the filopodial extensions. Filopodial extensions have been reported to target GABAergic neurons ^37,103^. Hence, we did not include extraction of filopodia in this work. The method does not apply deep learning-based techniques and can be scaled up to larger datasets with reasonable computational resources.

To segment individual vesicles from electron microscope (EM) images, we applied the deep-learning-based cell segmentation tool called Cellpose^67^. Cellpose provides a pre-trained model for detecting cell boundaries, trained on extensive image datasets. Direct deployment of this model for segmenting individual vesicles was initially unsuccessful due to differences in scale and morphology between cells and vesicles. Given that the cell diameter parameter significantly influences the segmentation accuracy, we enlarged the EM images using cubic interpolation via the resize function provided by Python’s OpenCV (cv2) package, such that synaptic vesicles have diameters around 20 pixels. Then, we fine-tuned the pre-trained Cellpose cytoplasm model and trained one additional model from scratch with 817 manually labeled synaptic vesicles (Extended Data Fig. 6e). The Cellpose software supported straightforward re-training through the PyTorch backend. We adopted a learning rate of 0.001 with a default decay strategy and momentum as implemented in Cellpose. The network could achieve the desired performance with 2,000 epochs of training on one NVIDIA GTX 1050 Ti GPU in about 2 hours. We used another manually labeled 535 vesicles as a test set, and the accuracy of the segmentation was evaluated using precision, recall and F1-score, treating each vesicle segment as a distinct object. Objects smaller than 100 pixels were treated as dust and removed from accuracy measurements. Neuron segmentation was used as a mask to filter out any vesicle segments falling outside the neuron boundaries.

### Identification of subcellular target locations of inhibitory axons

CAVE software including CaveClient, MeshParty, and Pcg_skel were used^39^. Given an inhibitory axon, we first picked out all of its output synapses targeting pyramidal cells whose soma existed within the image volume. For each identified postsynaptic pyramidal cell, we generated an approximate skeleton representation. This skeletonization was accomplished by employing the *pcg_skel.pcg_skeleton* function, where each level 2 chunk of the neuron was represented as a node and connected by edges to other level 2 chunks. The *root_point* parameter was set to the centroid of the pyramidal cell nucleus. To refine the skeleton for sufficiently high resolution, we applied linear interpolation of the skeleton nodes. The *meshparty.skeleton_io.export_to_swc* function was used, specifying a *resample_spacing* parameter to 1000 and setting *interp_kind* to ‘linear’ to perform the interpolation. We then mapped each output synapse of the inhibitory axon to the nearest node of the refined skeleton based on Euclidean distance. To measure how far each synapse was located from the postsynaptic pyramidal cell body, we converted the skeleton into an undirected graph and used the *networkx.dijkstra* function to compute the path length from each synapse-mapped node along the skeleton edges to the centroid of the pyramidal cell nucleus.

### Analysis of MF-Pyr connectivity

The MF-Pyr connectivity was analyzed as a bipartite graph from MF to pyramidal cells. A total of 636 Pyr cells and all of their inputs (∼25,200 MFs) in the dataset were used to construct a bipartite graph, wherein only connections with three or more synapses were included. To analyze MF-Pyr connectivity, we compared the reconstructed connectivity against two different null models, including a configuration model and an anatomically constrained model. In the MF-Pyr connectivity, MF out-degree is the number of target pyramidal cells, and the pyramidal in-degree is the number of presynaptic MFs. The configuration model preserves both in- and out-degrees but randomizes the partners. In the anatomically constrained random model, each bouton was randomly connected to the dendritic spines (thorny excrescences, TE) within a given radius (10 µm and 50 µm). MF boutons and pyramidal cell TEs were represented by the centroids of the MF-Pyr synapse coordinates. We computed pairwise Euclidean distances between all MF boutons and TEs. Temporary edges were assigned between bouton-TE pairs only if their separation was within the radius, ensuring zero connection probability beyond this radius. Within the radius, the probability of connection was uniform. The list containing these temporary edges was shuffled to randomize their order, and then each edge was evaluated sequentially. An edge was accepted only if neither the bouton nor the TE had already been connected by a previously accepted edge. Following this initial iteration, a subsequent iteration was performed to ensure that every bouton was associated with an accepted edge, preserving all MF boutons to have connections. For the configuration and radius model, 100 randomized models were sampled. We quantified the occurrence of all bipartite motifs involving three and four neurons (Fig. 4d). The 3-neuron motifs include fan-in and fan-out structures. The 4-neuron motifs include fan-in, fan-out, path-of-length-3, and path-of-length-4 (also known as the butterfly motif). Clustering coefficient *C* was computed on the projected graph of the MF-Pyr bipartite network, wherein each node represents a pyramidal cell and an edge between a pair of pyramidal cells (i.e. nodes) indicate they share input from a MF. In such a projected graph, high clustering means if Pyr-a shared inputs with Pyr-b, and Pyr-b shared inputs with Pyr-c, then Pyr-a and Pyr-c are more likely to share inputs.

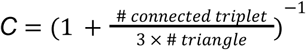

where a connected triplet is a set of three nodes where at least two edges connect them in a path (i.e., a center node connected to two others). A triangle is a set of three nodes where each node is connected to the other two. The factor of three accounts for the fact that each triangle contains three triplets.

We use participation ratio^56,60^ to measure the distribution of the network encoding capacity, based on reconstructed or null model connectivity.

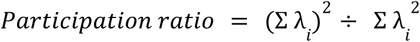

where λ are the eigenvalues of the covariance matrix for the synaptic weights. This participation ratio reflects dimensionality based on the connectivity ^56,60^.

## Data availability

The EM image data (rendered at 18 nm/pixel) and segmentation can be accessed at https://pyr.ai

A list of all identified cell types, along with links to the segmentation and EM data, can be found at https://codex.flywire.ai/research/mouse_ca3_explorer. An account is required, but access is open to all users.

All data, including the volumetric data and meshes, can be programmatically accessed through CAVE (https://github.com/CAVEconnectome) and cloudvolume (https://github.com/seung-lab/cloud-volume).

## Code availability

The analysis code is available at https://github.com/seung-lab/ca3_paper.git

The CA3 dataset uses CAVE for hosting proofreading and analysis (https://github.com/ CAVEconnectome).

## Acknowledgements

D.W.T and H.S.S. acknowledge support from the National Institutes of Health (NIH) BRAIN Initiative (U19 NS132720, RF1 MH123400) and CONNECTS programs (UM1 NS132250, UM1 NS132253), from S10 instrumentation program (S10OD023602), from the Simons Foundation, from the Princeton Neuroscience Institute, as well as assistance from Google and Amazon. Z.Z. acknowledges support from NIH BRAIN Initiative (K99 NS135650). The authors thank N. Kemnitz, D. Ih, K. Lee, T. Nguyen, J. A. Bae, J. Strout, A. Halageri, S. Popovych, T. Macrina of Zetta.ai for their reconstruction services; J. Wiggins, G. McGrath and D. Barlieb for computer system administration; R. Morey for data storage and website maintenance; M. Husseini for project administration; J. Maitin-Shepard for Neuroglancer; P. Schlegel, G. S. X. E. Jefferis for assisting with registration of the EM volume to the reference brain and processing cell type information; the management at SixEleven for coordination and proofreader management; J. Hebditch, R. Willie, K. Willie, C. David, and the SixEleven team for proofreading; W. A. Lee and J. Phelps for advice on image stitching; S. Dorkenwald, C. M. Schneider-Mizell, D. Brittain, and K. Wiley for general CAVE maintenance and troubleshooting support, which benefited all users, including users of this dataset; citizen scientists A. Kristiansen, A. Hernandez, A. Morren, J. Skelton, K. Kruk, M. Lichtenberger, N. Serafetinidis, R. Margossian, T. Stocks for proofreading; C. David, D. Jones for preparing training materials and community management for citizen scientists; K. Kuehner for assisting with website design; A. Matsliah for uploading cell type information to Codex. F.C. thanks the Allen Institute for Brain Science founder, P. G. Allen, for his vision, encouragement and support. The authors acknowledge the use of the Imaging and Analysis Center (IAC) operated by the Princeton Materials Institute at Princeton University, which is supported in part by the Princeton Center for Complex Materials (PCCM), a National Science Foundation (NSF) Materials Research Science and Engineering Center (MRSEC; DMR-2011750). The funders had no role in study design, data collection and analysis, decision to publish or preparation of the manuscript. The US Government is authorized to reproduce and distribute reprints for Governmental purposes notwithstanding any copyright annotation thereon.

## Author contributions

Z.Z.: conceptualization, sample preparation, serial sectioning, x-ray μCT imaging, electron microscopy imaging, image montaging, ground-truths, segmentation evaluation, proofreading management, cell typing, discoveries, analysis, figure preparation, and writing. C.P.: bouton and vesicle segmentation, anatomical measurements (including proximodistal and superficial-deep positioning, axon lengths, laminar organization, and inhibitory axon targeting of compartments), mossy fiber identifications, analysis, and figure preparation. E.W.H.: mossy fiber identifications and quantifications, cell typing, spine density quantification, proofreading quantification, segmentation evaluation, analysis, and figure preparation. R.L.: Segmentation. S.Y., M.S.: proofreading training and management for 611 team and citizen scientists. B.S.: ground-truth annotations (cell morphology, semantic, and synapse), mossy fiber identifications, proofreading, segmentation evaluation. C.S.J.: website design and creation, implementation of Spelunker neuroglancer, CAVE troubleshooting and maintenance. A.R.S.: website design and rendering, training materials, community management. W.M.S.: software for data transfer, flat segmentation, skeletonization, segmentation voxel statistics, fixing organelle segmentation holes. F.C.: onboarding, troubleshooting, and maintenance of CAVE infrastructure, synapse import, and predefined SQL query. H.S.S., D.W.T.: conceptualization, projection leadership, and writing.

---

**Extended Data Fig.1:**
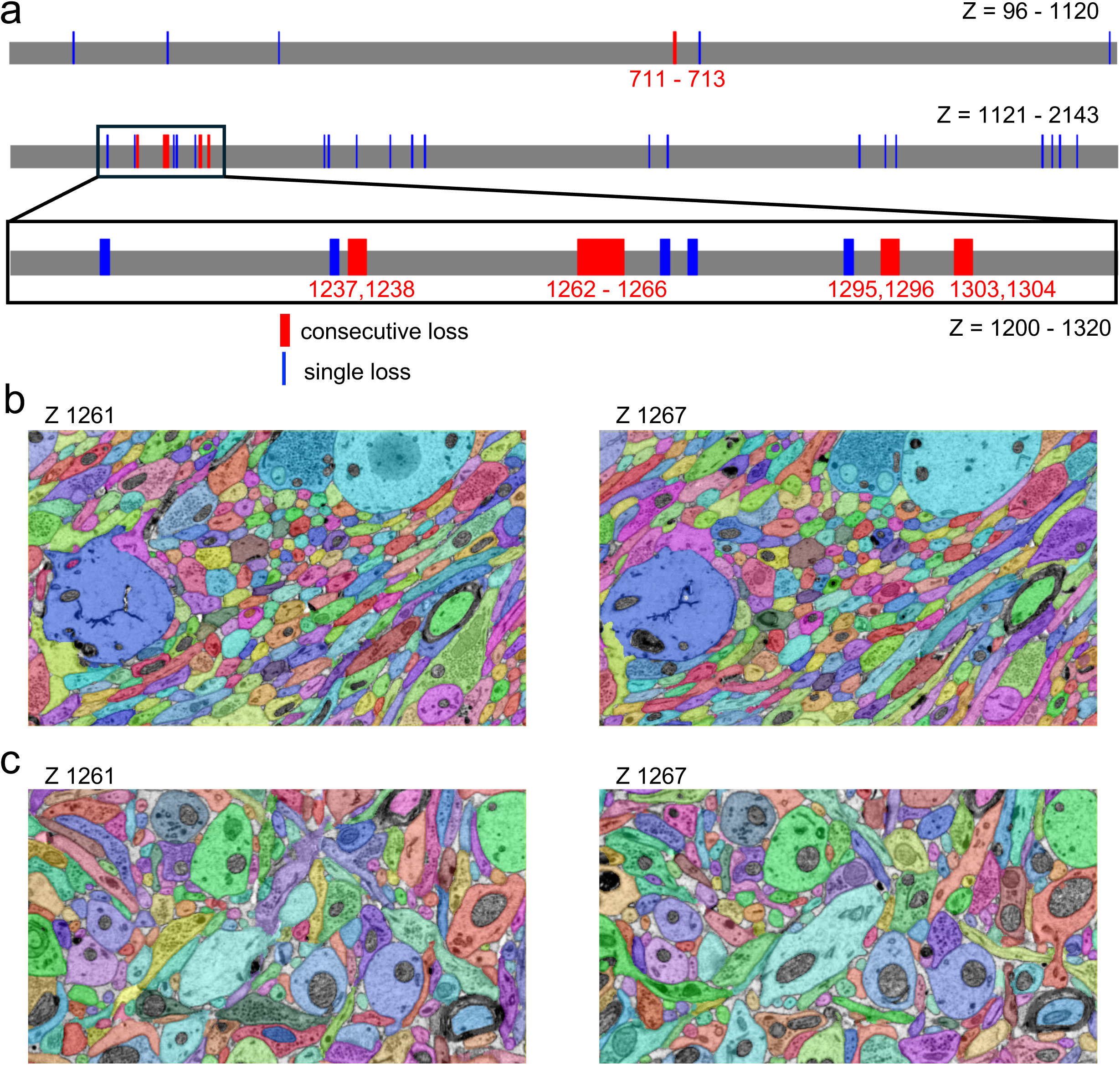
Dataset section map. **a,** Section loss is visualized across the segmented volume spanning z = 96 to z = 2143 (first and second rows). A region with concentrated consecutive missing sections is magnified in the third row. Red number indicates consecutive missing sectioning. **b - c**, Two regions of interest (ROIs) were exhaustively proofread across sections z = 1261 and z = 1267. ROI (b) contains 322 segments, primarily axons, while ROI (c) contains 162 segments, consisting of thin axons, dendrites, and spines. We identified probable continuations across the gap for all segments except for a small number of segments that are spine heads extending from dendrites and likely terminating shortly within the gap.

**Extended Data Fig.2:**
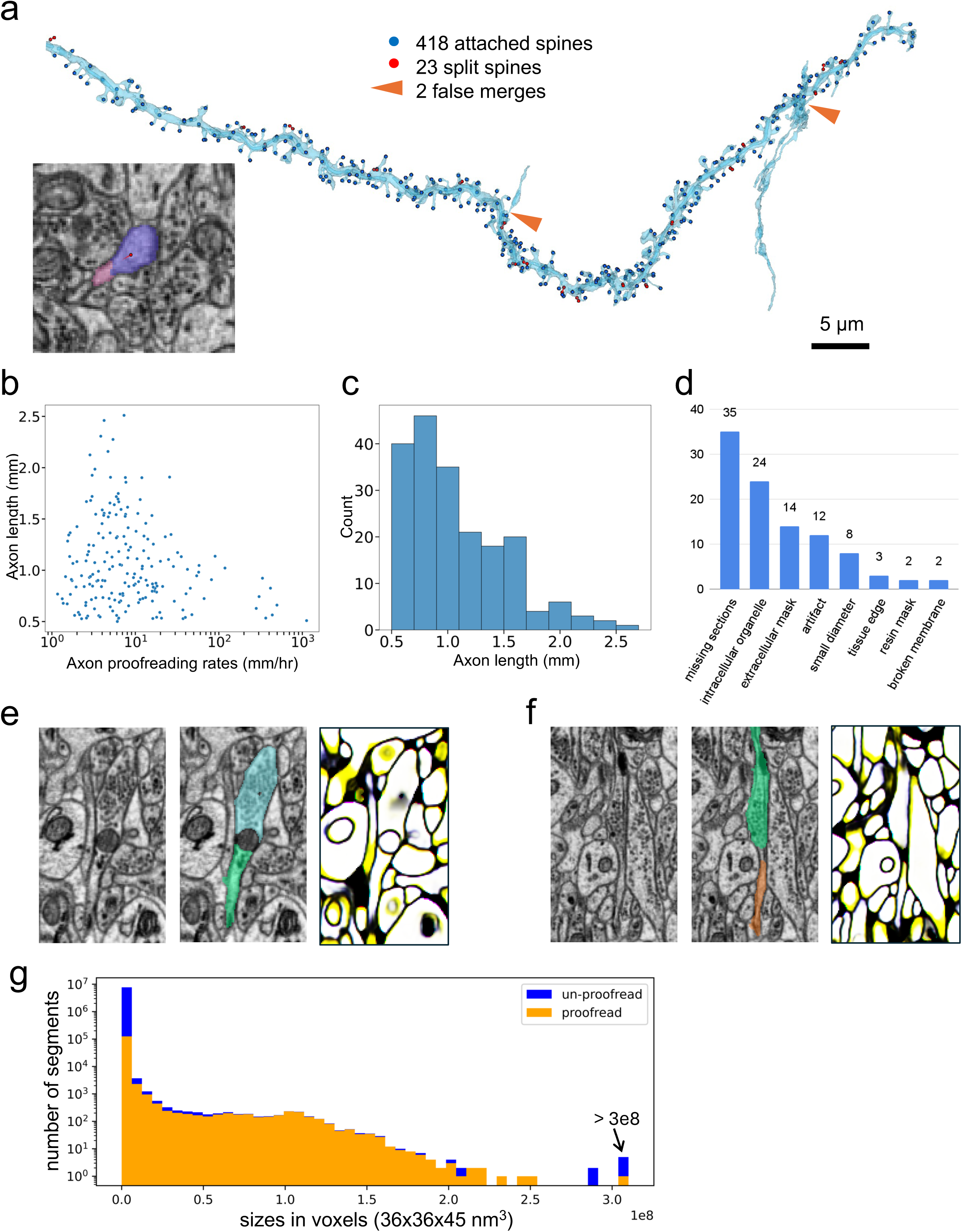
Segmentation quality. **a,** Assessment of spine attachment rate. Ground-truth spines were annotated in un-proofread segmentation, shown in blue. The automated segmentation achieved a recall of 94.8% and a precision of 99.5% for spine attachment. Inset shows an example of a segmentation split at a thin spine neck. **b - c,** Axon proofreading rates and axon lengths for pyramidal cells. Only pyramidal cells with axons longer than 0.5 mm were included in the calculation of proofreading rates. **d,** Categorization of the causes of 100 randomly selected segmentation errors. Artifacts include folds, cracks, and contaminations. **e,** A segmentation split error caused by an intracellular organelle, typically a mitochondrion, where the mitochondrial membrane merges with the cellular membrane. Shown are: EM image, EM image overlaid with the split segmentation, and 3D affinity map. The x, y, and z affinities are mapped to the red, green, and blue (RGB) channels, respectively. Low z-affinity combined with high x- and y-affinities appears as yellow (red + green). **f**, A segmentation split error due to incorrect extracellular masking. The small region between the green and orange segments shows very low affinity, resulting from incorrect labeling by a model trained to detect extracellular space. **g**, A histogram of segment sizes, measured in number of voxels (voxel size: 36 × 36 × 45 nm³), including only segments larger than 5,000 voxels. About 32% of voxels in the volume have been proofread.

**Extended Data Fig.3:**
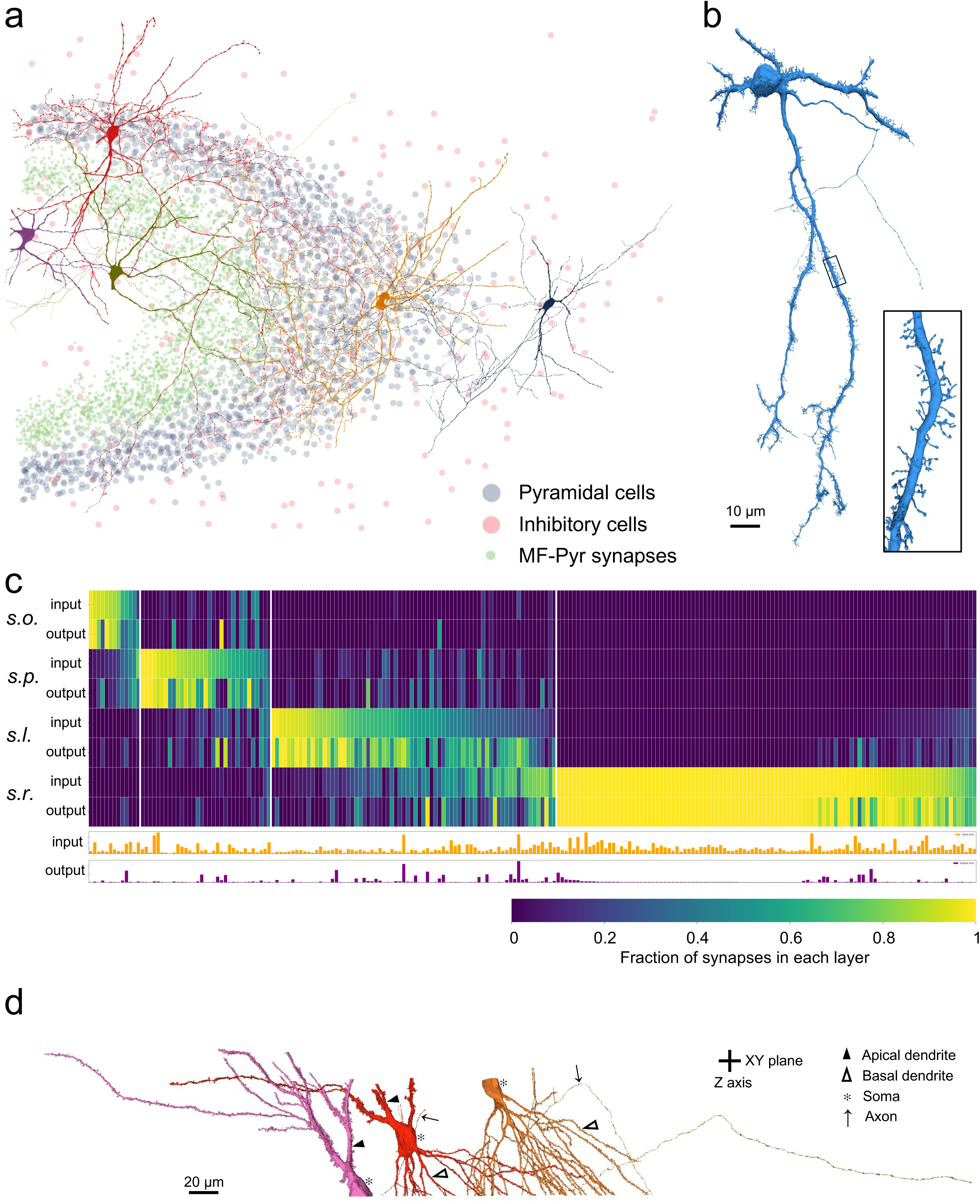
Classification of inhibitory cells. **a**, Visualization of inhibitory cells innervating different layers: stratum oriens (dark blue), stratum pyramidale (red), stratum lucidum (green), and stratum radiatum (purple). **b**, An inhibitory cell with spiny dendrites located in stratum lucidum. Spines on inhibitory cells tend to be longer, thinner, and more sparsely distributed compared to those on pyramidal cells. **c**, Fractions of input and output synapses by layer for inhibitory cells. Top: fraction of synapses per layer; bottom: total number of input and output synapses. s.o., stratum oriens; s.p., stratum pyramidale; s.l., stratum lucidum; s.r., stratum radiatum. **d**, Side view of reconstructed pyramidal cells, illustrating cases where cells are truncated at different compartments (basal dendrites, soma apical dendrites) due to limited z-axis.

**Extended Data Fig.4:**
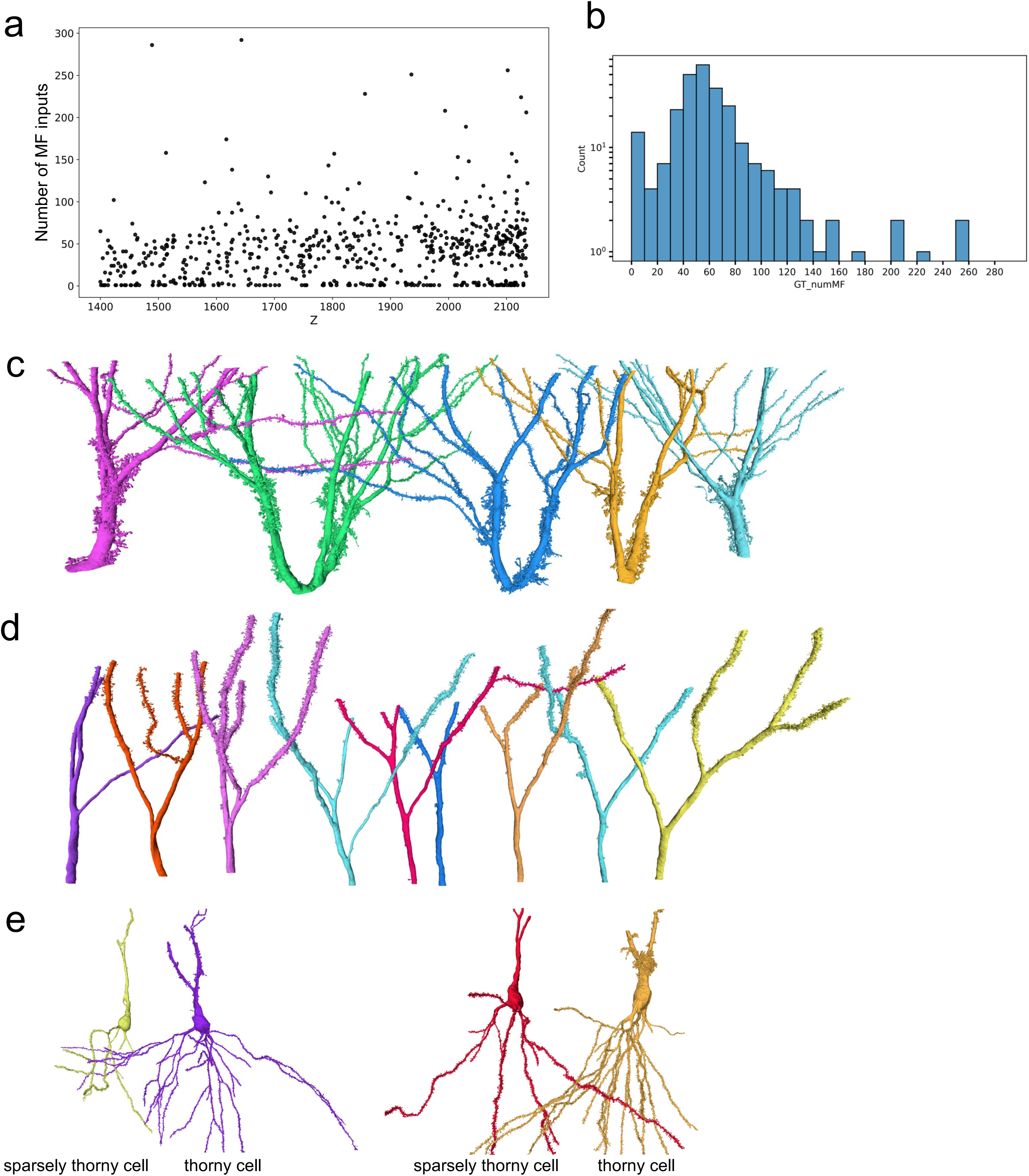
Mossy fiber inputs and morphology of CA3 pyramidal cells. **a,** Number of MF inputs for pyramidal cells at different Z levels. **b**, Distribution of number of MF inputs for a subset of long-apical pyramidal cells (266 cells) that have apical dendrites with side branches (e.g. those in Fig. 2c). **c - d,** Upper apical dendrites of representative thorny (b) and sparsely thorny (c) CA3 cells, shown truncated at the proximal apical dendrite, with their cell bodies located outside the volume. **e,** The basal dendrites of thorny cells are more elaborate than those of sparsely thorny cells.

**Extended Data Fig. 5:**
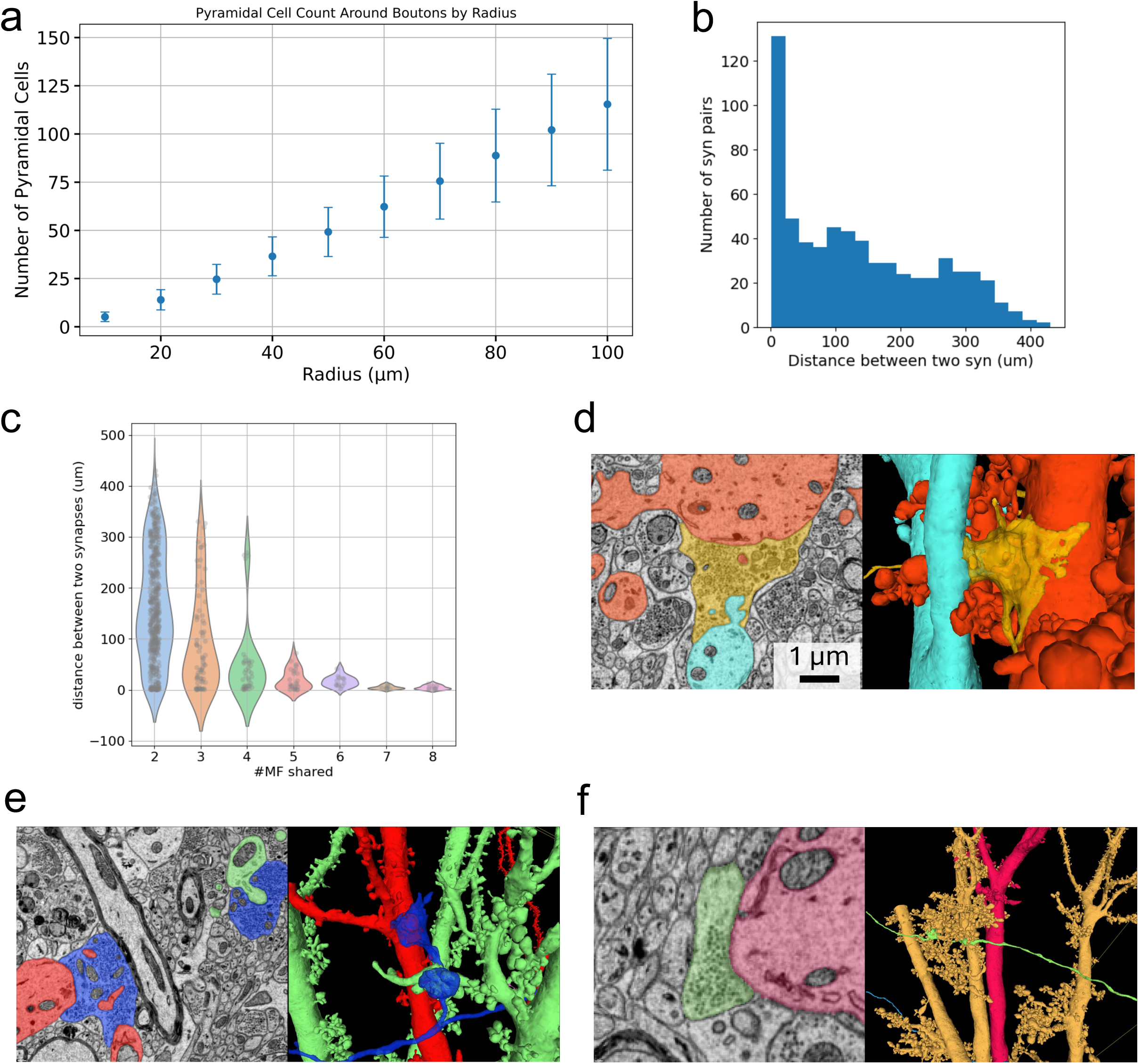
Pyramidal cells share mossy fiber inputs. **a,** The number of pyramidal cells (y-axis) was quantified based on the presence of thorny excrescence spines located within specified radii (x-axis) surrounding each mossy fiber (MF) bouton. **b-c,** Distances between synapses of pyramidal cell pairs that share two or more common MF inputs. **d,** A single MF bouton (orange) provides input to thorny spines of two neighboring pyramidal cells. **e**, A pair of neighboring MF boutons (blue) from a single mossy fiber provides input to two adjacent pyramidal cells, respectively. **f**, A synapse formed by a mossy fiber *en passant* bouton onto the dendritic shaft of a pyramidal cell (red).

**Extended Data Fig.6:**
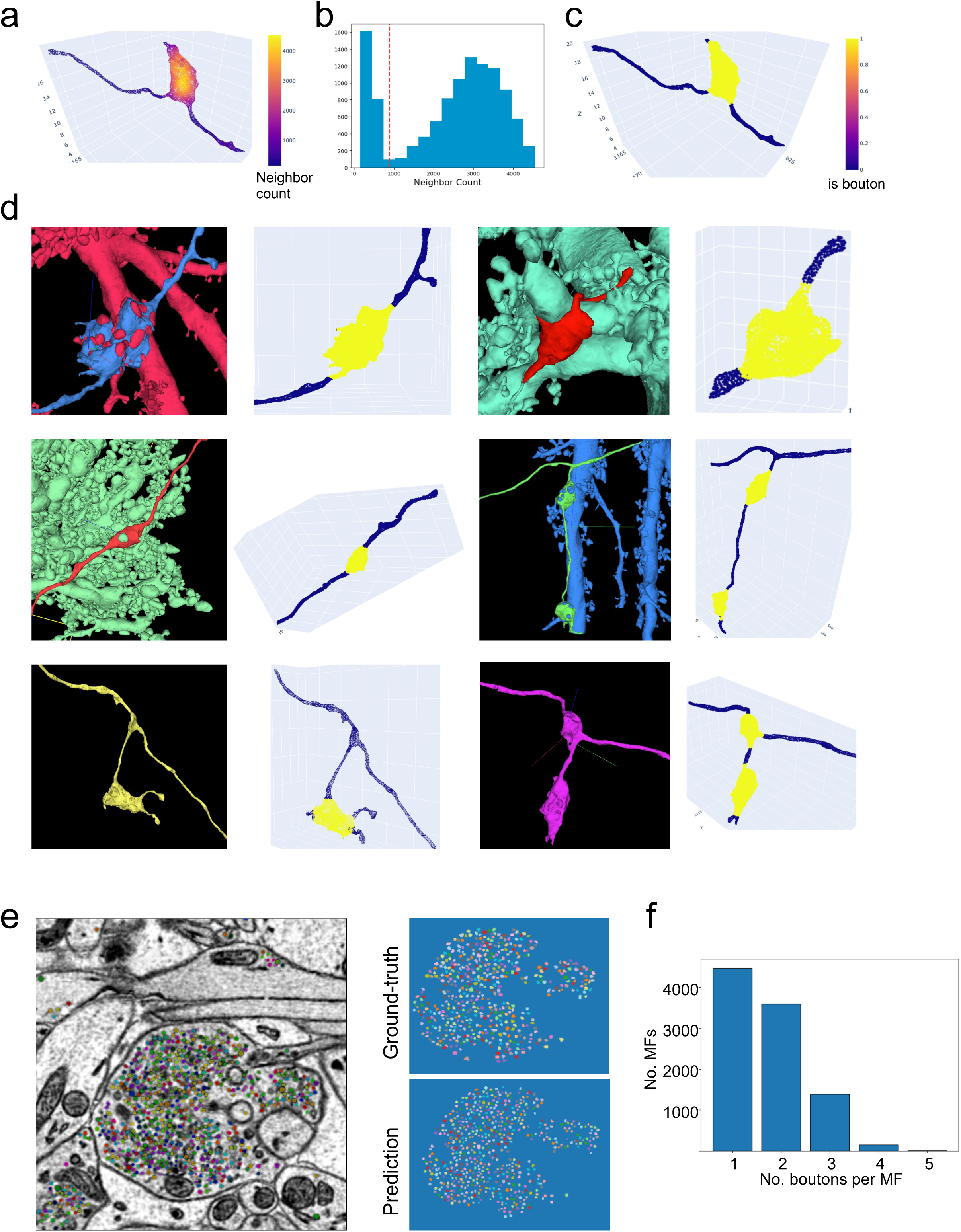
Automated segmentation of mossy fiber boutons and vesicles. **a,** A representative MF bouton extracted using a bounding box and color-coded by local mesh density, quantified as the number of mesh vertices within a 2.5 μm radius (neighbor count). **b,** Histogram of neighbor counts exhibits a bimodal distribution, corresponding to bouton and non-bouton regions of the 3D mesh. **c,** Voxel-wise probability of being a bouton based on a decision boundary derived from the distribution in a and b. **d,** Application of the segmentation method across diverse MF terminal morphologies, including large, terminal, small, doublet, boutons with filopodia, and doublets with short inter-bouton distances. Color scale as in c. **e,** Vesicle detection within an MF bouton using a model re-trained from a cellular segmentation method ^55^. Model performance on unseen data yielded a precision of 0.861, recall of 0.959, and F1 score of 0.907. **f**, Number of boutons per mossy fiber (MF) within the volume, quantified for MFs that form synapses with the 636 reconstructed pyramidal cells.

**Extended Data Fig.7:**
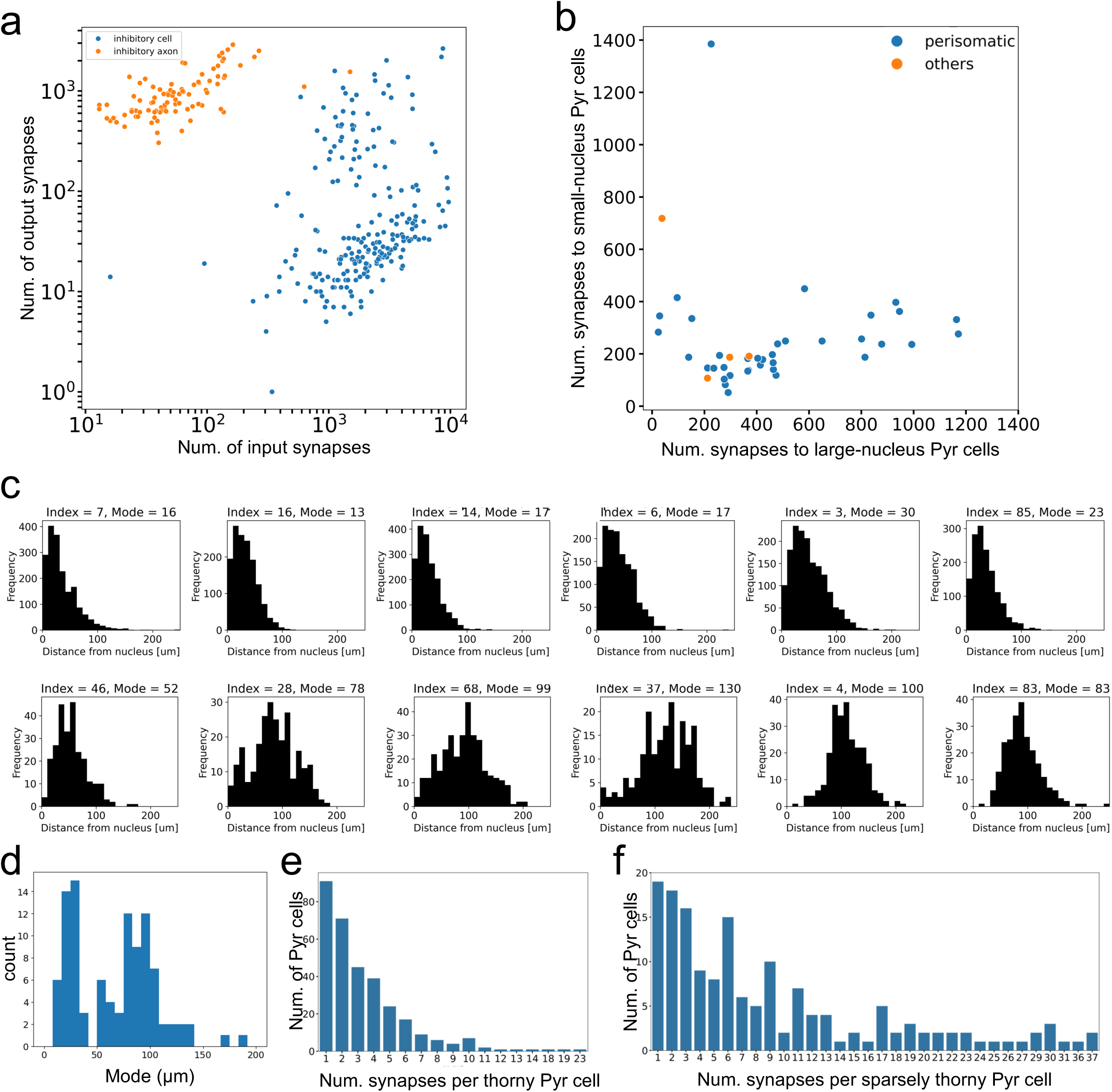
Perisomatic and dendritic targeting inhibitory cells. **a,** Number of input and output synapses for inhibitory cells and axons. Inhibitory axons have substantial arbors and output synapses within the volume, despite their somata lying outside the imaged volume. **b,** Number of synapses from individual inhibitory neurons onto pyramidal cells grouped by nucleus volume (threshold, 1232 µm^3^). Each circle represents the number of synapses from a single inhibitory cell. **c**, Distributions of along-the-skeleton distances from inhibitory synapses to the somata of postsynaptic pyramidal cells. Top: five perisomatic-targeting inhibitory cells with synapses located near the pyramidal soma (mode < 50 µm). Bottom: five dendrite-targeting inhibitory cells with most synapses located farther from the soma (mode > 50 µm). **d**, The modes for inhibitory cells exhibit a bimodal distribution. **e-f,** Synapse counts onto individual thorny and sparsely thorny pyramidal cells by a representative thorny-targeting and sparsely thorny-targeting inhibitory neuron, respectively.

**Extended Data Fig.8:**
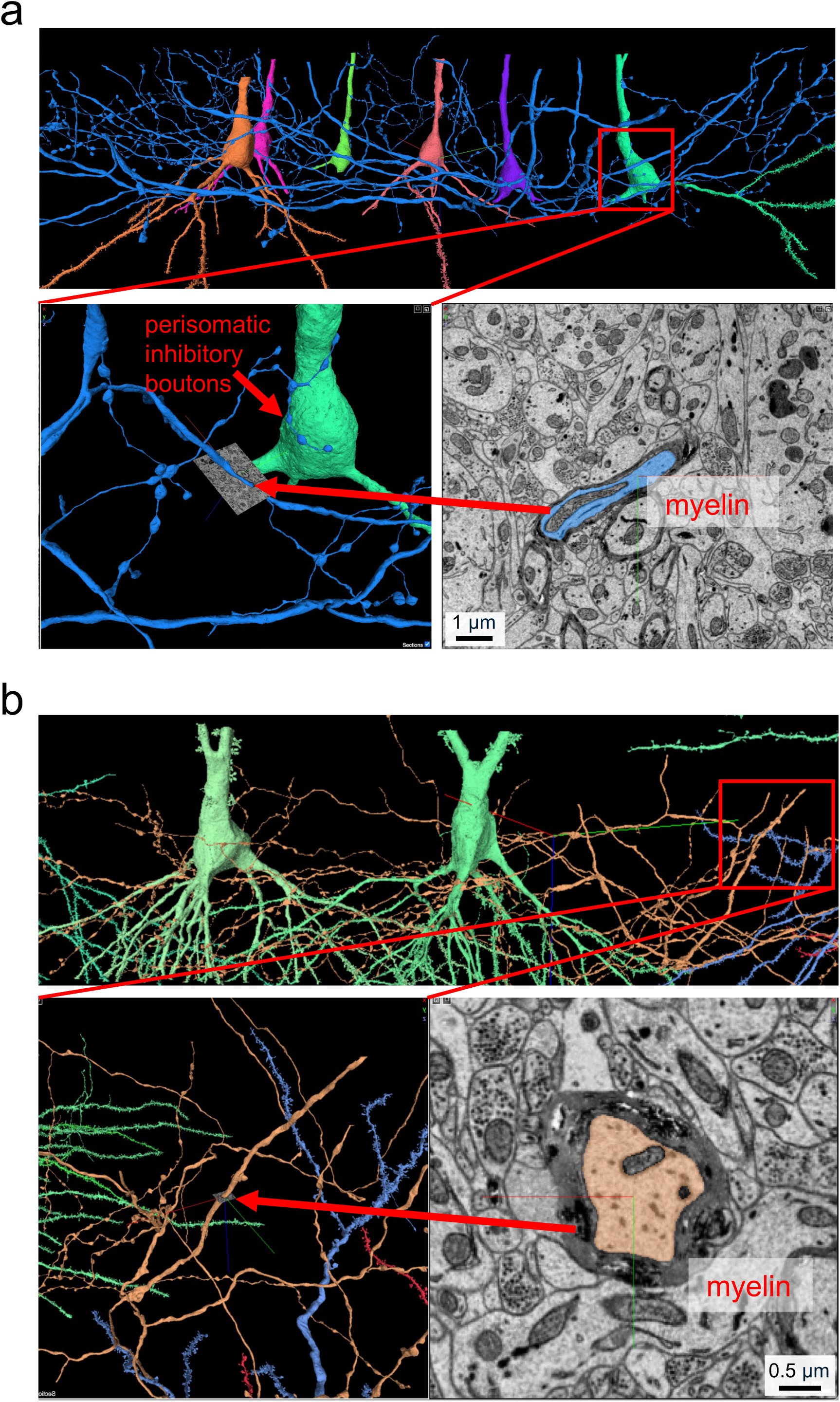
Sparsely thorny- and thorny-preferring perisomatic inhibitory axons are myelinated.

**Extended Data Fig.9:**
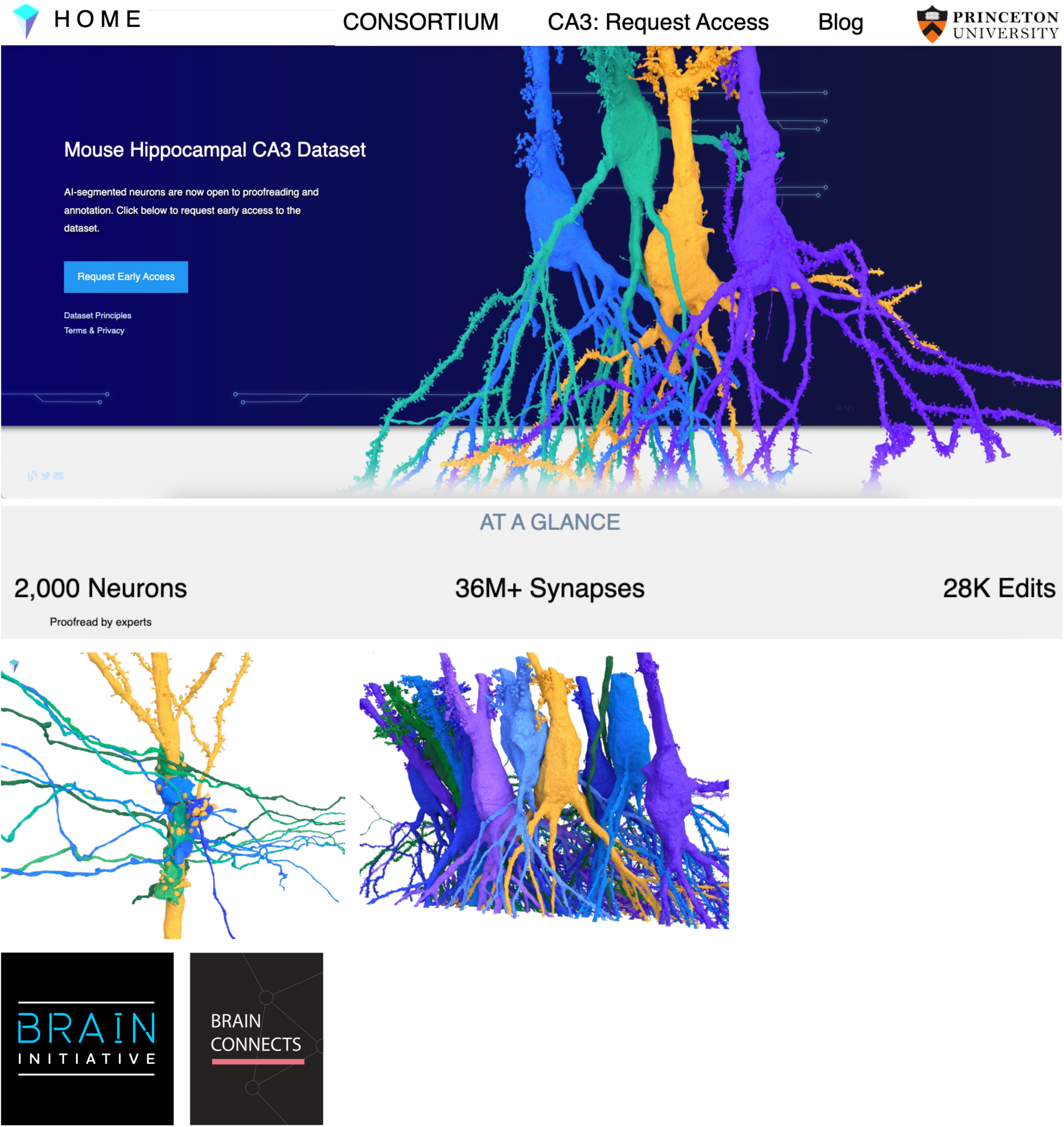
Pyr.ai, a platform for hippocampal connectomics.

